# Data-driven discovery of innate immunomodulators via machine learning-guided high throughput screening

**DOI:** 10.1101/2023.06.26.546393

**Authors:** Yifeng Tang, Jeremiah Y. Kim, Carman KM IP, Azadeh Bahmani, Qing Chen, Matthew G. Rosenberger, Aaron P. Esser-Kahn, Andrew L. Ferguson

## Abstract

The innate immune response is vital for the success of prophylactic vaccines and immunotherapies. Control of signaling in innate immune pathways can improve prophylactic vaccines by inhibiting unfavorable systemic inflammation and immunotherapies by enhancing immune stimulation. In this work, we developed a machine learning-enabled active learning pipeline to guide *in vitro* experimental screening and discovery of small molecule immunomodulators that improve immune responses by altering the signaling activity of innate immune responses stimulated by traditional pattern recognition receptor agonists. Molecules were tested by *in vitro* high throughput screening (HTS) where we measured modulation of the nuclear factor *κ*-light-chain-enhancer of activated B-cells (NF-*κ*B) and the interferon regulatory factors (IRF) pathways. These data were used to train data-driven predictive models linking molecular structure to modulation of the NF-*κ*B and IRF responses using deep representational learning, Gaussian process regression, and Bayesian optimization. By interleaving successive rounds of model training and *in vitro* HTS, we performed an active learning-guided traversal of a 139,998 molecule library. After sampling only *∼*2% of the library, we discovered viable molecules with unprecedented immunomodulatory capacity, including those capable of suppressing NF-*κ*B activity by up to 15-fold, elevating NF-*κ*B activity by up to 5-fold, and elevating IRF activity by up to 6-fold. We extracted chemical design rules identifying particular chemical fragments as principal drivers of specific immunomodulation behaviors. We validated the immunomodulatory effect of a subset of our top candidates by measuring cytokine release profiles. Of these, one molecule induced a 3-fold enhancement in IFN-*β* production when delivered with a cyclic di-nucleotide stimulator of interferon genes (STING) agonist. In sum, our machine learning-enabled screening approach presents an efficient immunomodulator discovery pipeline that has furnished a library of novel small molecules with a strong capacity to enhance or suppress innate immune signaling pathways to shape and improve prophylactic vaccination and immunotherapies.

## 1 Introduction

The efficacy of prophylactic vaccination and immunotherapeutics is predicated on effective stimulation of innate immune responses. In vaccination, for example, helper molecules known as adjuvants are often required to stimulate innate pathways involved in antigen presentation and processing that are critical in invoking a productive adaptive immune response. Despite the necessity of such signaling events to maximize the potency, excessive activation of signaling pathways by adjuvants can cause undesirable systemic inflammation, and limit tolerability and dosage in a clinical setting. ^1–3^ Conversely in immunotherapy, it is often essential to have strong stimulation to improve the immunogenicity and mitigate suppression from a tumor micro-environment.^4,5^

Two major effectors of the innate immune response are the nuclear factor *κ*-light-chain-enhancer of activated B-cells (NF-*κ*B) pathway and the interferon regulatory factors (IRF) pathway. The NF-*κ*B pathway plays an essential role in inflammation as well as immune activation, while the IRF pathway produces type-I interferons that are essential for a productive antiviral response. ^6–8^ Among these signaling pathways, pattern recognition receptors (PRRs) are cellular receptors expressed on immune cells that identify pathogen-associated molecular patterns (PAMPs) and initiate a cascading immune response. PRRs are necessary for the activation of antigen-presenting cells (APCs) that act as a link between the innate and adaptive immune responses and play a critical role in detecting and responding to pathogens.^9,10^ PRR agonists are molecules that bind to PRRs, mimicking the effects of pathogenic molecules and triggering an immune response. As such, PRR agonists have recently been used as adjuvants to activate both NF-*κ*B and IRF pathways and are the most common targets for manipulating the innate immune response.^11^ A breadth of PRR agonist-based adjuvants comprised of pathogenic motifs have been used as adjuvants in vaccines and immunotherapies, such as lipopolysaccharides (LPS)^12^ and Monophosphoryl Lipid A (MPLA)^13^ that target toll-like receptor (TLR) 4, synthetic oligodeoxynucleotides that contain unmethylated cytosine-phosphate-guanine dinucleotide motifs (CpG-ODN) targeting TLR9,^14^ cyclic guanosine monophosphate-adenosine monophosphate (cyclic GMP-AMP, cGAMP) that binds and activates the stimulator of interferon genes (STING), ^15^ polyriboisosinic:polyribocytidylic acid [poly(I:C)] that are recognized by TLR3, ^16^ an imidazoquinoline synthetic small-molecule R848 that targets TLR7/8,^17^ and flagellin that activates TLR5.^18^ Although these adjuvants can be potent activators of immune responses, a well-known limitation of current popular adjuvants is excessive and uncontrolled inflammation.^1,11,19^ This has motivated efforts to discover novel adjuvants with reduced inflammation profiles,^20,21^ but it has proved challenging to develop novel PRR agonists capable of specifically tuning the level of stimulation in inflammatory pathways without disrupting the desired stimulation along immune activation pathways.

An alternative approach to regulating the innate immune response is through immunomodulators – molecules co-delivered with PRR agonists to reduce inflammation or otherwise modulate innate immune stimulation by enhancing or suppressing innate immune signaling pathways. Moser *et al.*^20^ have demonstrated that a selective NF-*κ*B inhibitor known as SN50 has such immunomodulation capacity. SN50 is a cell permeable peptide that consists of nuclear localization sequence of the NF-*κ*B subunits p50 and blocks the import of p50-containing dimers into the nucleus.^22,23^ It was found that SN50 can reduce the levels of inflammatory cytokines TNF-*α* and IL-6 while enhancing antigen-specific antibody titers when delivered with the TLR9 agonist CpG.^20^ As compared to peptides like SN50, small molecules present attractive candidates for immunomodulators due to their better synthetic accessibility and reduced potential for immunogenicity. To this end, Moser *et al.*^21^ also demonstrated that a small molecule NF-*κ*B inhibitor honokiol and its derivatives can be used as immunomodulators with similar functions to SN50. More recently, we conducted a targeted experimental screen of a small molecule library of *∼*3000 compounds, many of which were known to influence the immune system, to discover, after removing cytotoxic compounds using the same viability filter as applied in this study, novel molecules capable of suppressing NF-*κ*B activity by up to 9-fold, elevating NF-*κ*B activity by up to 7-fold and elevating IRF activity by up to 7-fold. ^24^

The hypothesis underpinning the present work is that immunomodulators of greater potency and diversity may be discovered by screening large libraries of small molecules. A challenge in extending experimental screening efforts in this manner is the vast size of molecular space: the number of pharmacologically active molecules obeying the Lipinski rules has been estimated to be in excess of 10^60^.^25,26^ (To help place this number in context, it is estimated that there are “only” 10^22^ stars in the visible universe.^27^) Experimental screening can only probe a small fraction of these molecules due to time, labor, and materials constraints. Human intuition and experience present valuable heuristics to guide this search, but the relative infancy of immunomodulator discovery efforts, absence of detailed mechanistic understanding, and vast size of molecular space can make these heuristics limiting and subject to human preconceptions, bias, and potential blind spots. Data-driven models trained in concert with experimental screens offer a systematic means to guide traversal of molecular design space using predictive models of immunomodulation activity learned on-the-fly from the screening data. Such models, often referred to as quantitative structure activity relationship (QSAR) or quantitative structure property relationship (QSPR) models, ^28–30^ have been employed in multiple applications to molecular design and discovery, including self-assembling *π*-conjugated peptides,^31^ cardiolipin-selective small molecules, ^32^ battery electrolytes,^33^ energy storage materials,^34^ and nanoporous materials. ^35^ QSAR models have a venerable history dating back to the development of cheminformatics and combinatorial chemistry in the 1980’s^36^ and have recently benefitted from the integration of powerful generative deep learning tools. ^37–39^

This work reports our development of a data-driven pipeline integrating machine learning and *in vitro* high throughput screening (HTS) to accelerate the discovery of small molecule immunomodulators of the innate immune response. We curated a library of 139,998 candidate small molecules from commercial chemical screening libraries readily available for purchase. These chemical screening libraries are pre-filtered to be structurally-diverse and drug-like. Commencing with limited experimental measurements, we constructed a data-driven QSAR model integrating deep representational learning,^37,40^ Gaussian process regression,^41^ and Bayesian optimization^42^ to guide subsequent rounds of molecular selection and experimental HTS. After screening only 2,880 compounds comprising *∼*2% of the molecular search space using the QSAR/HTS framework, we discovered nine molecules with an unprecedented ability to either up- or downregulate the activity of NF-*κ*B or IRF when activated by known agonists. The most successful candidates from our study demonstrated remarkable improvements in modulating immune activity compared to those identified in our earlier screen. ^24^ Considering the magnitude of the immunomodulation response relative to that induced by specific agonists, these top-performing candidates realized improvement of up to 110% in elevating NF-*κ*B activity, 83% in elevating IRF activity, and 128% in suppressing NF-*κ*B activity. Additionally, we identified 167 novel immunomodulators with at least 2-fold enhancement or suppression over transcription factor activity of interest, which represents a 105% increase in the total number of known immunomodulators with this level of activity, and nine novel immunomodulators with at least 10-fold activity modulation.

In addition to specialists – immunomodulators capable of enhancing or suppressing an immune signaling pathway when delivered in concert with a specific agonist – we also discovered a number of generalists – immunomodulators capable of modulating immune signaling pathways when co-delivered with a number of agonists. These generalists, although less effective in immunomodulation for any specific agonist, represent promising versatile agents to incorporate into diverse prophylactic vaccines and immunotherapies due to their broad spectrum compatibility.

Finally, we conducted additional characterization assays of 17 top-performing candidates identified in our screen to measure their cytokine release profiles in primary cells. These molecules demonstrated significant capacity to modulate the secretion of various cytokines, including upregulating TNF-*α* production by over 10-fold, downregulating TNF-*α* production by over 16-fold, and upregulating IFN-*β* production by over 6-fold. One of our top candidates demonstrated a 3-fold enhancement in IFN-*β* production when delivered with cGAMP, which is a strong stimulator of interferon genes (STING) agonist.

A deficiency of the VAE/GPR models is that despite their predictive power in guiding the HTS campaign, the design rules mediating the mapping from small molecule chemical structure to immunomodulatory profile is not readily available from these relatively “black box” models. Accordingly, we performed a *post hoc* analysis of the molecules considered in our screen using interpretable “glass box” linear regression models that are expected to possess lower predictive accuracy than the nonlinear VAE/GPR models but can transparently identify particular chemical fragments that are principal drivers of specific immunomodulation behaviors. We found that the presence of halogen moiety in immunomodulators is highly predicative of suppression of NF-*κ*B activity regardless of the type of agonist. The carbonyl and carboxyl moiety is predictive of suppression of the immune responses in NF-*κ*B pathway activated by TLR4 agonists such as LPS and MPLA, and predictive of enhancement of the immune responses in the NF-*κ*B pathway activated by CpG. It was also discovered that aromatic heteroatom moieties are predictive of enhancement in NF-*κ*B activity and of suppression in IRF activity. Even though IRF pathway suppression is of limited clinical interest, this information could be used to guide the development of IRF enhancing immunomodulators by either avoiding the inclusion of aromatic heteroatom moieties or removing such chemical fragments to enhance their potency. This analysis offers design rules to modify the structure of immunomodulators for achieving practical immunomodulation goals in vaccine design or immunotherapy development, enriching candidate libraries with molecules predicted to have promising immunomodulatory behaviors.

The data-driven QSAR computation detailed in this work presents a powerful framework to integrate with experimental HTS to construct on-the-fly models of immunomodulatory activity from the screening data and use these models to efficiently traverse molecular candidate space and extract interpretable design rules. We also observe that this integrated computational and experimental screening platform detailed herein is quite generic, and can be easily repurposed to other early-stage molecular discovery pipelines to guide efficient search and optimization within large molecular spaces by modular context-specific replacement of the molecular library and experimental assays.

## 2 Experimental Methods

### 2.1 High-throughput screening assays

We chose transcription factor levels of NF-*κ*B and IRF activity as our two measures of immunomodulator performance when delivered in combination with one of four PRR agonists: lipopolysaccharides (LPS),^12^ Monophosphoryl-Lipid A (MPLA),^13^ CpG ODN 1826 (CpG),^14^ and 3’3’-cGAMP (cGAMP).^15^ NF-*κ*B transcription levels are correlated with inflammatory responses and should typically be minimized to reduce reactogenicity in prophylactic vaccinations. Under certain circumstances, enhanced NF-*κ*B activity is instead required to optimize the efficacy of vaccines.^7,8^ IRF activity, which is related to interferon production, should typically be maximized for a productive antiviral response. ^8,43^ Following previously established protocols, we employed RAW-Dual™ macrophages as a reporter cell line that can quantitatively report NF-*κ*B and IRF activity via secreted alkaline phosphatase (SEAP) and Lucia luciferase using the proprietary substrates QUANTI-Blue™ and QUANTI-Luc™ for absorbance and luminescence readings.^24^ We seeded 50,000 RAW-Dual™ macrophages in 384-well plates in 45 *µ*L of complete media using a MultiDrop™ Combi liquid handler, then incubated for 1 hour at 37*^◦^*C. We transferred immunomodulator compounds from source plates (10mM in DMSO) to a final concentration of 10 *µ*M using a Janus^®^ G3 via pintool. One column and two rows at the edge of each plate were left blank and filled with water to avoid possible systematic errors due to evaporation. Following 1 hour incubation at 37*^◦^*C, one of the four PRR agonists was added in 5 *µ*L of media to achieve the desired concentration. The concentration of each agonist is reported in the Supporting Information. Cells were incubated with or without agonist overnight until transcription factor activity was analyzed. This activity was measured via absorbance at 620 nm or luminescence readings using a BioTek Synergy™ Neo2 Hybrid multimode microplate reader. In the modulator-only negative control, we were able to measure the net enhancement or suppression in activity of the screened compounds on the cells in the absence of agonists. By employing a supernatant based assay, we were able to simultaneously observe the NF-*κ*B and IRF activity within a single well. The raw readings of absorbance and luminescence of each well were then divided by the average reading of the positive controls (presence of agonists and absence of immunomodulators) which are on the same plate to define the fold change associated with each modulator relative to the baseline of the corresponding agonist. This plate-based normalization ensures measurement consistency by eliminating plate-to-plate and day-to-day variance. Each immunomodulator was incubated in two replicated plates and the results were averaged. Experimental errors are calculated from the standard deviation of the mean calculated from the two replicates with the same immunomodulator using standard propagation of errors. A schematic illustration of the experimental screening process is presented in Figure S10. An illustration of the plate layout used in the HTS experiments is presented in Figure S11.

The small molecule library source plates were hosted by the University of Chicago Cellular Screening Center (Chicago, Illinois, USA) and purchased from various vendors. The RAW-Dual™ cells, QUANTI-Blue™, QUANTI-Luc™ and the PRR agonists were purchased from InvivoGen (San Diego, California, USA). A comma-separated values (CSV) file providing SMILES strings of all compounds along with their vendor and results of our active learning screen is provided in the Supporting Information.

### 2.2 Viability filter analysis

As some modulators may be cytostatic or cytotoxic to cells, viability was monitored after overnight addition by monitoring confluency via IncuCyte^®^ imaging.^44,45^ Two confluency masks were generated using IncuCyte^®^ software over all imaged wells, and modulators were determined non-viable if both sets of confluency masks were lower than 70% of those of resting cells. This methodology was validated with select library plates using a traditional Promega^®^ CellTiter-Glo^®^ assay. Immunomodulators considered non-viable are not necessarily cytostatic or cytotoxic in other cell lines or clinical settings, so immunomodulator candidates exhibiting extraordinary behaviors were not categorically ruled out on the basis of this viability assay alone. However, unless explicitly stated otherwise, we only report on immunomodulators measured to be viable under this assay.

### 2.3 Measurement of cytokine release profiles in bone marrow derived dendritic cells (BMDCs)

A subset of the top performing candidate molecules identified in the high-throughput active learning screen were subjected to low-throughput measurement of cytokine release profiles in murine bone marrow derived dendritic cells (BMDCs). Monocytes were harvested from 6-week-old C57BL/6 mice and were differentiated into dendritic cells using supplemented culture medium: RPMI 1640 (Life Technologies, Waltham, Massachusetts, USA), 10% HIFBS (Sigma-Aldrich, Burlington, Massachusetts, USA), Recombinant Mouse GM-CSF (carrier-free; 20 ng/ml; BioLegend, San Diego, California, USA), 2 mM L-glutamine (Life Technologies), 1% antibiotic-antimycotic (Life Technologies), and 50 *µ*M *β*-mercaptoethanol (Sigma-Aldrich). After 6 days of culture, BMDCs were plated at 100,000 cells per well and incubated with modulator (10 *µ*M). After 1 hour, one of the three tested agonists (LPS, CpG, cGAMP) was added. Cells were incubated for 24 hours at 37*^◦^*C and 5% CO_2_. Supernatant cytokines were measured using LEGENDplex™ Mouse Inflammation Cytokine Kit (Biolegends) or a VeriKine™ IFN-*β* ELISA (PBL Assay Science, Piscataway, New Jersey, USA). The cytokines measured by LEGENDplex™ are TNF-*α*, IFN-*β*, IFN-*γ*, IL-6, IL-1*α*, IL-1*β*, IL-10, IL-12p70, IL-17A, IL-23, IL-27, MCP-1, and GM-CSF.

## 3 Computational Methods

We developed a data-driven active learning framework to guide experimental screening of immunomodulator candidates. An overview of our integrated screening approach is illustrated schematically in Figure 1. The machine learning component of the framework comprises a deep representational learned projection of all 139,998 molecular candidates into a smooth and low-dimensional latent space suitable for regression and optimization using a variational autoencoder (VAE; Figure 1B), the construction of learned QSAR models over this space using Gaussian process regression (GPR, Figure 1C), and the selection of new candidate molecules from the design space using Bayesian optimization (BO, Figure 1D). The selected candidate molecules are then passed to experimental HTS by automated robotics (Figure 1A). Experimental measurements are then used to retrain and update the GPR models and inform subsequent rounds of BO candidate selection within a virtuous cycle of QSAR model training and model-guided experimental screening. Multiple passes through the active learning loop are conducted until desired performance metrics are reached or we encounter a performance ceiling evinced by a lack of improvement over a particular number of cycles. We now proceed to describe the VAE, GPR, and BO components of the machine learning framework.

**Figure 1:**
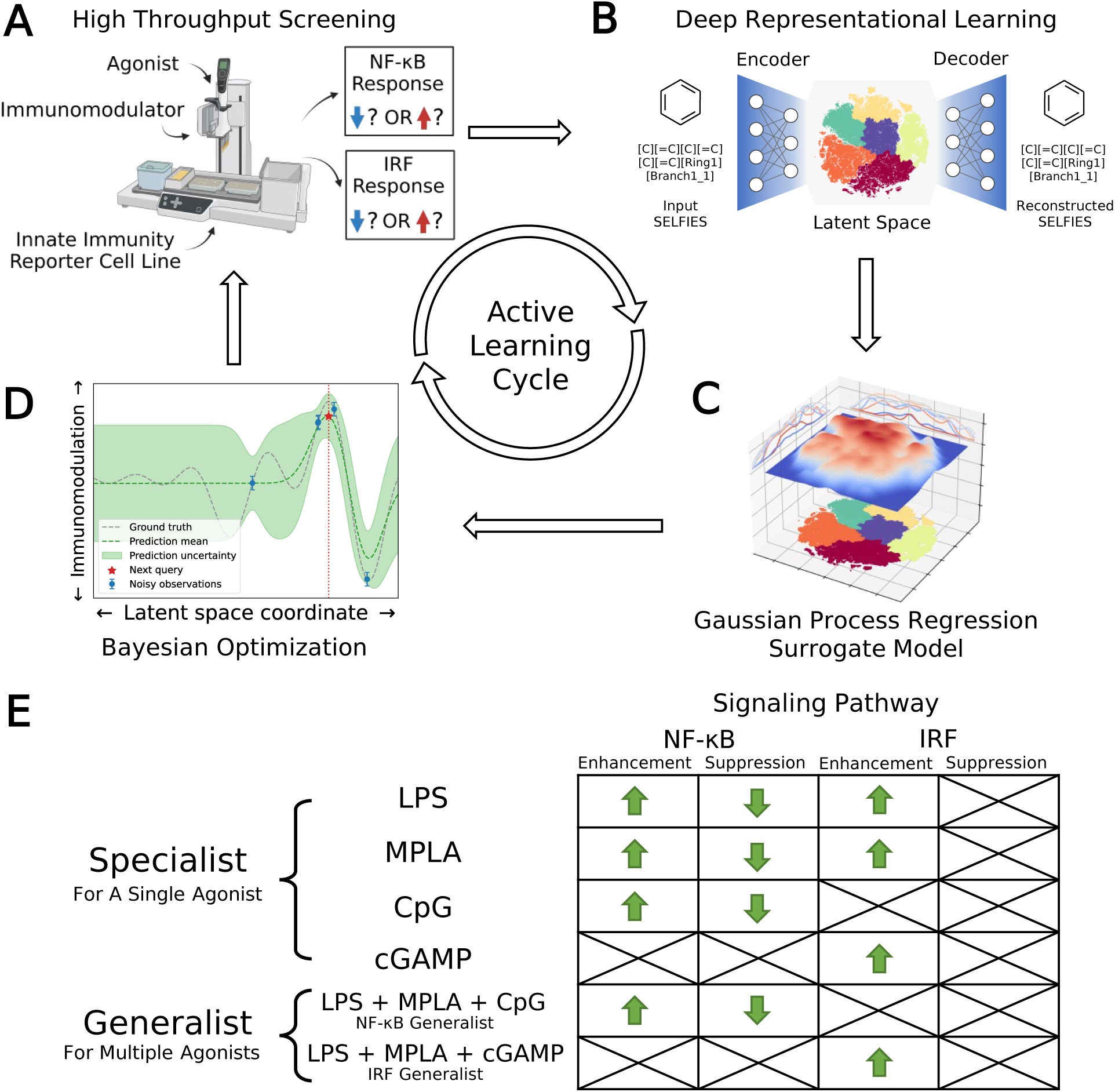
Data-driven active learning framework for immunomodulator discovery. (A) Immunomodulator candidates are subjected to *in vitro* high throughput screening (HTS) using automated liquid handling platforms. (B) Deep representational learning with a variational autoencoder (VAE) is used to learn an embedding of all 139,998 immunomodulator candidates into a smooth, low-dimensional latent space amenable to regression and optimization. (C) A supervised surrogate Gaussian process regression (GPR) model is trained to predict the immunomodulation of transcription factor levels in the NF-*κ*B and IRF pathways from all accumulated experimental measurements to date. (D) The trained surrogate model is interrogated using Bayesian optimization (BO) to select the most promising batch of as-yet-untested immunomodulators for the next round of experimental screening. Experimental measurements are then used to retrain and update the GPR models and inform subsequent rounds of BO candidate selection within a virtuous cycle of QSAR model training and model-guided experimental screening. (E) Optimization objectives comprising eight agonist-pathway combinations and twelve functional goals (12 green arrows). Upregulation and downregulation of the NF-*κ*B response are both important immunomodulation objectives, leading us to split these agonist-pathway combinations into two goals associated with enhancement and suppression. Only upregulation of the IRF response is a preferred objective, meaning that only enhancement of this response is targeted. We seek specialist immunomodulators capable of delivering large immunomodulatory effects when co-delivered with one particular agonist, and generalist immunomodulators capable of delivering large immunomodulatory effects when co-delivered with any one of a particular agonist group. Specifically, we seek an NF-*κ*B generalist capable of enhancement or suppression of the NF-*κ*B pathway when delivered in concert with any one of the LPS, MPLA, or CpG agonists, and an IRF generalist capable of enhancing the IRF pathway when delivered in concert with any one of the LPS, MPLA, or cGAMP agonists. Portions of this figure were created using BioRender.com.

### 3.1 Deep representational learning using variational autoencoders (VAEs)

Previous work has shown that variational auto-encoders (VAE) can be used to perform deep representational learning of discrete chemical structures to embed them into low-dimensional, continuous latent space embeddings well-suited to property regression and molecular optimization.^37,40,46^ The VAE comprises two consecutive deep networks: an encoder converting the discrete chemical representations into fixed-length vectors defining an embedding into a latent space, and a decoder that inverts this operation to reconstruct the discrete chemical representations from their latent space projections^40^ (Figure 1B). Chemical structures are represented to the VAE as SELFIES strings^46^ flattened into one-hot vectors. The loss function for network training comprises two objectives that are simultaneously optimized: accurate reconstruction of the chemical representations by decoding from their latent space projections and preservation of a prior distribution – frequently a multi-dimensional Gaussian – over the latent space. Conceptually, training of the network discovers a low-dimensional fixed-length vector representation of each molecule in the training data that preserves sufficient information to permit its accurate reconstruction within a compact distribution that promotes generalization to unseen data and from which it is easy to sample. As such, the latent space embedding furnishes a smooth, low-dimensional embedding in which chemical similarity of the training molecules is related to latent space proximity and that is well-suited to traversal and optimization by active learning (Section 3.2 and 3.3).

We constructed and trained the VAE model in PyTorch^47^ and its parameters were optimized under 5-fold cross-validation (CV). We achieve good performance employing a 500-200-100 fully-connected feedforward network architecture as the encoder, and a stack of two gated recurrent units (GRUs) as the decoder. An additional 100-node layer following the encoder network serves as the 100D latent space, and it is followed by the decoded network. We trained the VAE over an training library consisting of the 139,998 immunomodulator candidates (see CSV file in the Supporting Information) augmented with 1,108,666 molecules extracted from the ZINC library of commercially available small molecules ^48–51^ and other commercially available compound libraries for virtual screening. Although we consider only the 139,998 immunomodulator candidates within our screen, the relatively small size of this library means that it is beneficial to enrich the training set with additional small molecule candidates to train more stable models with improved generalizability and a smoother latent space. The VAE was trained only once at the beginning of the active learning search to produce a latent space embedding of all immunomodulator candidates that was held fixed throughout all subsequent iterations of the process. Training was conducted on a NVIDIA RTX 2080 GPU card requiring 9032 epochs and 1344 GPU-hours of training over the 1,248,664 candidate augmented training library. Full details of library definitions, model training, and hyperparameter optimization are provided in the Supporting Information.

### 3.2 Gaussian process regression (GPR) surrogate model training

The primary goals of our molecular screen are to discover novel immunomodulators to enhance or suppress the NF-*κ*B response as a means of, respectively, upregulating innate immune stimulation or downregulating inflammation in prophylactic vaccination to promote efficacy and safety, and to enhance the IRF response to upregulate production of type-I interferons and promote antiviral responses in cancer immunotherapies. This motivated us to discover molecules capable of enhancing the NF-*κ*B response, suppressing the NF-*κ*B response, and enhancing the IRF response. We did not search for molecules to inhibit IRF pathway, since this is presently of limited clinical significance.

We consider four agonists in this work: LPS, MPLA, CpG, and cGAMP. LPS and MPLA both target TLR4, CpG targets TLR9, and cGAMP targets STING.^12–15^ We sought to discover molecules that are *specialists* in achieving large immunomodulatory effects when delivered with one particular agonist, and those that are *generalists* in doing so when delivered with any one of a particular group of agonists. For example, a molecule that enhances the NF-*κ*B response when delivered with LPS would be regarded as a specialist enhancer of NF-*κ*B operating through the TLR4 receptor, whereas one that suppresses the NF-*κ*B response when delivered with any one of LPS, MPLA, or CpG would be regarded as a generalist suppressor operating through the TLR4 and TLR9 receptors. We expect that generalists may not be as good at immunomodulation in concert with any single agonist, but offer better immunomodulation profiles across multiple agonists. We quantify the performance of a specialist for a particular agonist via the fold change in the NF-*κ*B or IRF transcription factor activity induced by co-delivery of the immunomodulator with the agonist relative to delivery of the agonist alone. We quantify the performance of a generalist over a group of agonists as the average fold-change over that group.

We illustrate in Figure 1E the eight agonist-pathway combinations representing the 12 functional immunomodulatory goals of our immunomodulator screen. The rows of the matrix comprise the agonists or groups of agonists, and the columns comprise the signaling pathway and enhancement / suppression thereof. Considering the columns of the table, we recall that downregulation of the IRF pathway is of limited clinical interest and so is not included in our screening goals. Considering now the rows, we seek immunomodulatory specialists that exert large enhancement or suppressive effects when co-delivered with LPS, MPLA, CpG, or cGAMP agonists alone. We also seek an NF-*κ*B generalist capable of enhancement or suppression of the NF-*κ*B pathway when delivered in concert with any one of the LPS, MPLA, or CpG agonists, and an IRF generalist capable of enhancing the IRF pathway when delivered in concert with any one of the LPS, MPLA, or cGAMP agonists. We note that the cGAMP agonist is known to primarily affect the IRF pathway and so was not considered as either a specialist or generalist for NF-*κ*B immunomodulation. Similarly, CpG is known to primarily affect the NF-*κ*B pathway, and so was not considered as either a specialist or generalist for IRF immunomodulation.

Having defined our 12 functional goals it was the next task to optimize these functions over the smooth, low-dimensional latent space embedding of the 139,998 immunomodulator candidates learned by the VAE.^37^ To do so, we trained 12 independent GPR surrogate models employing Gaussian (a.k.a. radial basis function) kernels to learn an empirical mapping from the coordinates of each molecule within the 100D latent space embedding to each of the 12 objective functions (Figure 1C). In each round of the active learning screen, the GPR models were trained over all immunomodulator candidates for which we had experimental measurements of the level of modulation of the NF-*κ*B and IRF responses, and were then used to predict the performance of all remaining candidates for which experiments had not yet been performed. This is the key step in our data-driven screening process – the trained GPR models enable us to interpolate/extrapolate from the experimentally measured performance of a small number of candidates to predict the performance of all unmeasured candidates before actually conducting the experimental measurements. In this way, the surrogate models guide a prospective traversal of the candidate space by allowing us to focus the time, labor, and expense of experimentation toward the most promising molecules. Importantly, the performance predictions of the GPR models in each of the 12 design goals are also equipped with uncertainty estimates. As such, we can account for the typically higher model uncertainties when making extrapolative predictions to molecules that lie far away in the latent space (i.e., are more chemically dissimilar) from those that have already been measured. In the first round of our active learning screen, initial GPR models were trained over measurements for 2674 molecules within the candidate space for which we previously conducted experimental screening in our prior work.^24^ In subsequent passes through the active learning loop we retrained the GPR models over these original measurements plus all new measurements conducted in subsequent passes through the loop. Full details of the GPR kernel, training, and hyperparameter tuning are provided in the Supporting Information.

### 3.3 Multi-objective Bayesian optimization (BO) candidate selection

The GPR models were then interfaced with a multi-objective Bayesian optimization (BO) framework to select the best compound candidates for experimental testing in the next round of active learning^35^ (Figure 1D). For each of the 12 GPR models, we made performance predictions on all as-yet-untested molecules and scored each one according to the Expected Improvement (EI) acquisition function.^52^ The EI acquisition function accounts for both the mean and uncertainty of the GPR predictions to balancing exploitation and exploration to identify candidates most likely to lead to improved performance. To select molecules across the 12 performance goals, we integrated the 12 GPR models with a multi-objective Kriging believer batched sampling protocol to define a batch of 720 molecules for experimental testing.^53,54^ Specifically, we collated from each of the 12 GPR models the molecule that had not previously been selected for testing with the largest acquisition function value. This group of 12 molecules, with duplicates removed, was then used to retrain all 12 GPR models under a Kriging believer approach and the GPR models polled again for their next top-ranked molecules. We repeated this process until a batch of at least 720 molecules were selected and sent for experimental testing. Importantly, all selected molecules, regardless of their source GPR models, were subjected to experimental testing to evaluate their immunomodulatory profiles across all agonist types in both the NF-*κ*B and IRF pathways. Hence, molecules selected by one GPR model were not only used to retrain that particular GPR model, but also used to retrain all other models. Full details of the BO acquisition function, integrated Kriging believer batched sampling and stopping criteria analysis are provided in the Supporting Information.

### 3.4 Convergence assessment

One cycle of VAE embedding, GPR training, BO sequence selection, and experimental screening completes one loop of the active learning cycle (Figure 1A-D). We use two methods to monitor and determine convergence of the active learning loop. First, we employ a stabilizing predictions method to evaluate stabilization of the specialist GPR model predictions.^55^ To do so, we set aside a randomly selected 100,000 candidate stop set and measure the average Bhattacharyya distance^56^ *D_B_* between the GPR posterior evaluated over this stop set in successive rounds of the active learning screen. Large average *D_B_* values indicate that the GPR posterior is still being updated over the course of additional screening rounds and thus the convergence has not been reached yet, whereas small values indicate that additional rounds are not changing the GPR predictions and can be seen as an indicator for convergence. Second, we employ the performance difference method to assess the specialist GPR predictive performance by conducting 5-fold cross-validation over the accumulated labeled samples (i.e., candidates for which experimental assay measurements are available).^57^ When the absolute value of the cross-validated mean average error (MAE) on the labeled data reaches an acceptably low level and/or plateaus over the course of successive rounds, this indicates that the predictive power of the trained GPR is no longer changing with the accumulation of additional screening data and can be taken as an indication of model convergence. Generalist GPR models are not considered in this convergence assessment, because the immunomodulatory profiles of generalists are defined as linear combinations of specialists with different agonists in a particular group. Hence, the convergence of generalist GPR models is closely correlated with the convergence of specialist GPR models, and we choose to only assess the convergence of specialist GPR models. Using these criteria, we terminated the active learning screen after four rounds, during which we experimentally assayed a total of 2,880 compounds comprising *∼*2% of the 139,998-candidate molecular candidate space.

### 3.5 Inference of chemical design rules

At the conclusion of the active learning screen, we attempted to extract human interpretable design rules relating simple chemical properties of the immunomodulator candidates to their measured performance. To do so, we constructed simple “glass box” linear regression models linking the occurrence of particular structural fragments to the measured fold-change in the immunomodulatory response using least absolute shrinkage and selection operator (LASSO) regression.^58,59^ This resulted in sparse linear models with relatively few non-zero linear coefficients that identify those structural fragments that are the principal discriminants of the measured immunomodulatory activity.

To train these models, we combined the dataset consisting of 2880 molecules tested in this study with the 2674 molecules obtained in a prior study.^24^ We then eliminated any nonviable and redundant compounds to arrive at a dataset of 3560 distinct and viable molecular structures, along with their associated immunomodulatory profiles. We used the open-source cheminformatics software RDKit^60^ to featurize each of the 3560 experimentally assayed immunomodulators as a numerical vector of 85 substructure occurrences. We then eliminated eight irrelevant features that showed no variation across all the immunomodulators and seven redundant features that had a linear correlation *ρ*>0.95 with any other features. ^61^ This left us with *k* = 70 features for each immunomodulator that were compiled into the feature matrix *F ∈* ℝ^(*N*=3560)×(*k*=70)^. Each feature row comprising the feature values for each immunomodulator was normalized to unit length. A table detailing the selected substructure features, their chemical interpretation, and the feature selection process is provided in the Supporting Information.

Given the 3560*×*70 feature matrix *F*, we then constructed LASSO regression models to predict the activity change for each agonist-pathway combination: (A) NF-*κ*B–LPS, (B) NF-*κ*B–MPLA, (C) NF-*κ*B–CpG, (D) NF-*κ*B–Generalist, (E) IRF–LPS, (F) IRF–MPLA, (G) IRF–cGAMP and (H) IRF–Generalist (Figure 1E). Each LASSO regression model corresponding to an immunological objective is trained to predict the log2-fold change in immunomodulatory activity for an immunomodulator using the corresponding normalized feature vector. This training involves minimizing the L1 regularized loss, where the L1 penalization prevents overfitting by retaining only a small number of generalizable features present in our training dataset. The optimal number of features to use in our model is determined by 5-fold cross-validation on the L1 regularization weight. We identify the regularization weight values that result in the lowest generalization error, as well as the number of non-zero coefficients and mean absolute error (MAE) for predicting immunomodulation in log2-fold change corresponding to that optimal regularization weight. By examining the coefficients with the largest magnitudes in this optimal linear model, we can rank the molecular descriptors based on their immunomodulatory effect.

## 4 Results and Discussion

### 4.1 Active learning identifies novel candidate immunomodulators

We conducted four rounds of active learning-guided experimental screening of a library of 139,998 putative immunomodulators. In each round, we trained a GPR surrogate model over the experimental screening data collected to date as a surrogate predictor of immunomodulatory activity along 12 functional goals over eight agonist-pathway combinations. We then conducted BO to select a total of 720 candidates predicted to strongly enhance the NF-*κ*B response, inhibit the NF-*κ*B response, or enhance the IRF response in the presence of one particular agonist (i.e., a specialist) or in the presence of any one of a group of agonists (i.e., a generalist). We terminated our screen after four rounds, corresponding to a screening of 2880 immunomodulator candidates comprising *∼*2% of the molecular candidate search space.

In Figure 2A we quantify the extent of immunomodulation for each of the 12 functional goals by reporting the fold change in immune activation induced by the combination of PRR agonists and immunomodulators relative to that induced by agonists alone for the molecules considered in each round of the active learning screen. Importantly, we validate that immunomodulators alone do not stimulate an immune activation in the absence of agonists (Figure S6). The goal of the active learning screen was to perform on-the-fly learning of a QSAR model to guide the optimal selection of the most promising immunomodulator candidates and achieve round-on-round improvements in the identification of top performers. We observe the preponderance of molecules are clustered around a fold-change of unity, meaning that they have a very limited effect on immune activation, and it is the rarer molecules in the tails of the distributions that are of primary interest and to which active learning directs our screen. Looking at the most potent immunomodulator in different function goals (i.e., the maximum or minimum of the distribution in fold change illustrated as orange and purple bands in Figure 2A), we observe clear round-on-round improvements in 11/12 functional goals, indicating that the screen is resolving novel high-performing candidates. Only the LPS specialist to enhance the IRF response shows no significant improvement after the first round, perhaps indicative of the relative paucity of immunomodulators for these goals within the candidate space. Two functional objectives – MPLA specialists to enhance the NF-*κ*B response and MPLA specialists to enhance the IRF response – show a continuing upward trend after four rounds of screening, but all other functional goals appear to have reached a plateau.

**Figure 2:**
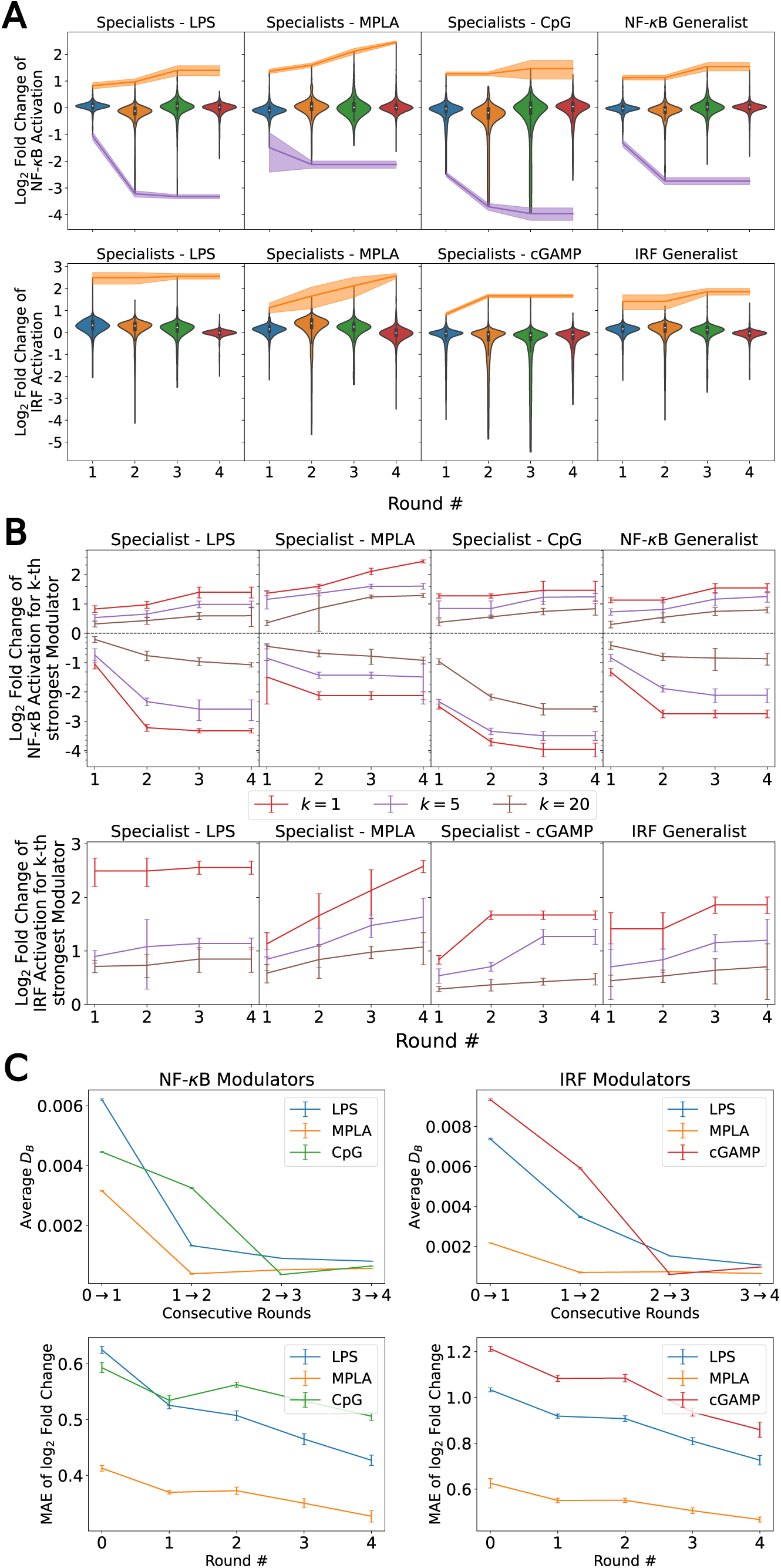
Results of the data-driven immunomodulator discovery active learning screen. (A) Violin plots of the active learning screen over the 12 functional objectives over the eight agonist-pathway combinations of interest. Fold changes in the NF-*κ*B and IRF responses are measured relative to delivery of the agonist alone. Each violin contains the 720 molecules experimentally assayed in each round along the eight agonist-pathway combinations. The orange and purple lines show, respectively, the top enhancer and inhibitor identified by the mean of the two independent experimental measurements, and the shading shows the range of the two measurements. We are most interested in molecules populating the tails of the distributions exhibiting strong ability to alter the immune response by enhancing the NF-*κ*B response, inhibiting the NF-*κ*B response, or enhancing the IRF response in the presence of one particular agonist (i.e., specialists) or in the presence of any one of a group of agonists (i.e., generalists). The NF-*κ*B generalist comprises the LPS, MPLA, and CpG agonists, whereas the IRF generalist comprises the LPS, MPLA and cGAMP agonists. (B) Fold changes in the *k*^th^ strongest enhancer/inhibitor in the 12 functional goals as a function of active learning round for *k* = 1 (red), *k* = 5 (purple), and *k* = 20 (brown). Data are reported as the mean of two independent measurements and error bars show the range of the two measurements. (C) Convergence assessment of the active learning screen for the specialist prediction goals. The Bhattacharyya distance *D_B_* between successive GPR posteriors over a randomly selected stop set of 100,000 points plateaus near a value of zero between Rounds 3 and 4 (upper row). Error bars report the standard error in *D_B_* estimated over the stop set. The 5-fold cross validated MAE in the predicted log2-fold change in immunomodulatory activity over all labeled data collected to date (lower row) shows a decreasing trend in all predictive goals indicating that additional predictive performance of the GPR may be gained under additional screening rounds.

In addition to discovering those top-performing immunomodulators, we also seek to expand the number of immunomodulators to the tails of the distributions to identify multiple novel high performing immunomodulators. In Figure 2B, in addition to the top-performing immunomodulators, we show the log2-fold change of the 5^th^ and the 20^th^ strongest enhancer and suppressor for each functional goal with respect to each round. Similarly, we observe round-on-round improvements in 11/12 functional goals for the 5^th^ and the 20^th^ strongest immunomodulator curve, again with the LPS specialist to enhance the IRF response being the only exception. This indicates that the screen is exposing high-profile immunomodulators to enrich the tails.

To quantify convergence of the GPR surrogate models, we employed the stabilizing predictions method by computing the average Bhattacharyya distance *D_B_*between GPR posteriors in successive rounds over a randomly selected stop set of 100,000 points and employed the performance difference method by computing the 5-fold cross-validated mean average error (MAE) over the accumulated labeled data collected to date as a function of screening round.^31,55–57^ As illustrated in Figure 2C, all specialist GPR models exhibited convergence in *D_B_* by Round 4. Figure 2C indicates that the models have not yet fully converged with respect to the MAE, indicating that the predictive capacity could be further improved by additional rounds of screening. However, given the plateau trends in the active learning screen (Figure 2B), the convergence of the average Bhattacharyya distance, and the expensive nature of an additional round of experimental screening, we elected to terminate the search after four rounds under the rationale that the marginal returns of additional screening rounds are likely to be small and that a large number of high-performing candidates in all 12 functional objectives have been discovered within the first four rounds. Nevertheless, it is possible that more performant molecules could be identified by additional rounds of screening.

We present in Figure 3 projections of the candidate molecules into a 2D t-distributed Stochastic Neighbor Embedding (t-SNE) compression of the 100D VAE latent space. We hesitate to accord too much interpretation to these 2D compressed representations, but do observe that the distribution of sampled points in the learned latent space embedding is consistent with the GPR/BO driving broad exploration of the molecular candidate space and has not become overly focused or stuck in any one particular region over the course of the active learning screen (Figure 3A-C). Immunomodulators with potent enhancement or suppression performance are distributed broadly within the 2D t-SNE embeddings (Figure 3D), although it appears that there may be some localization of high-performance suppressors in the top-left corner of the space.

**Figure 3:**
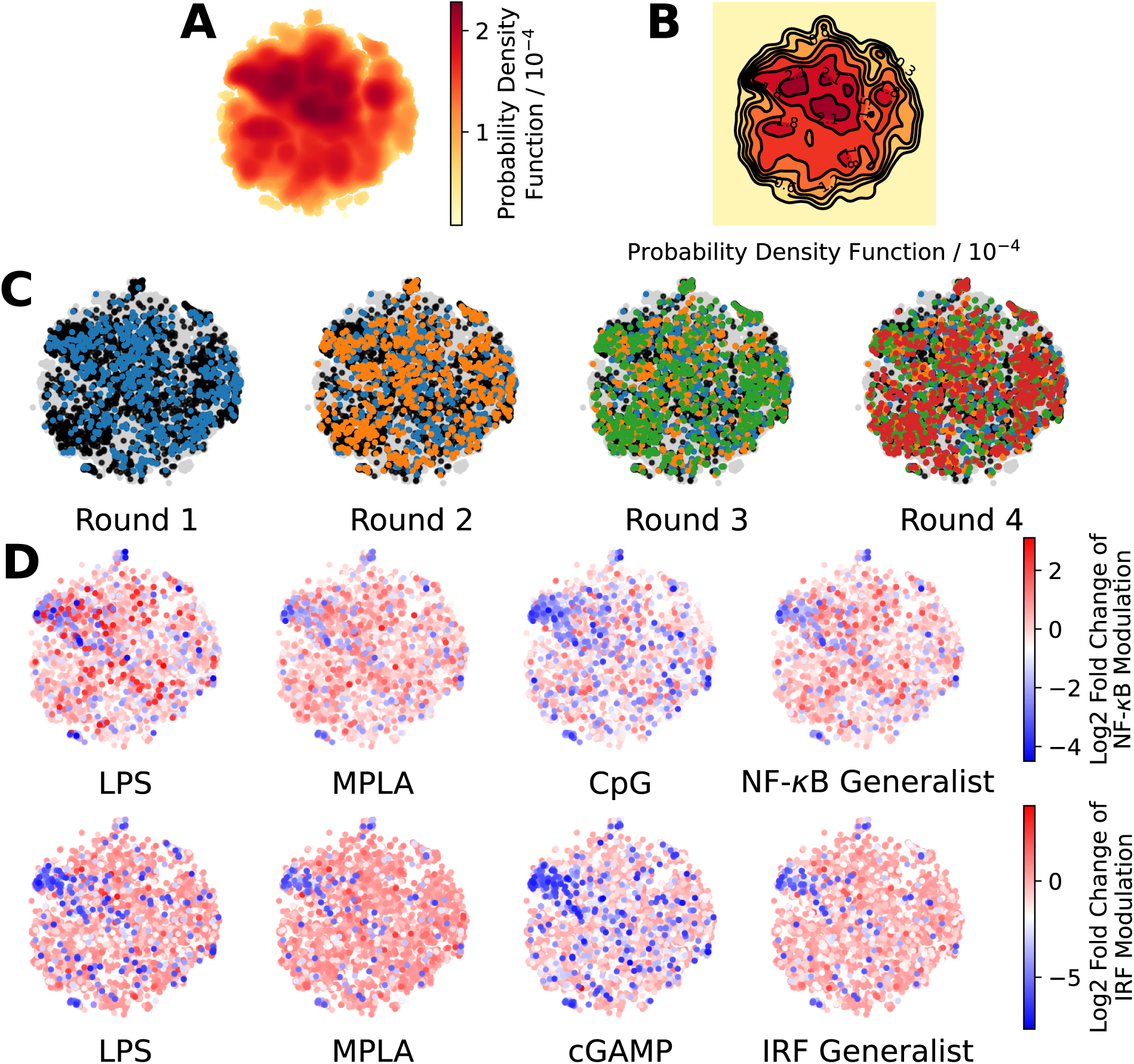
Measuring the progress of the active learning screen over 2D projections of the 100-dimensional latent space. (A) Probability density function estimated by kernel density estimation of a projection of the 142,672 molecules, consisting of 139,998 candidate molecules and 2,674 molecules from previous study^24^ into a 2D t-distributed Stochastic Neighbor Embedding (t-SNE) embedding of the 100D VAE latent space. (B) A contour plot of the 2D pdf presented in panel A. (C) Identification of the newly selected molecules in each of the four rounds of the active learning screen within the 2D t-SNE embedding: 139,998 molecules defining the complete candidate space (grey), 2674 molecules screened in prior work^24^ used to train the initial GPR models (black), 720 molecules selected in Round 1 (blue), 720 molecules selected in Round 2 (orange), 720 molecules selected in Round 3 (green), and 720 molecules selected in Round 4 (red). (D) Measured immunomodulatory effects of all molecules for which experimental assay measurements are available projected into the 2D t-SNE embeddings.

We can retrospectively estimate the savings in time and labor realized by the four-round active learning search in comparison to a naïve random sampling baseline. Although this is an intrinsically imperfect analysis since we do not know the immunomodulatory activity of the unmeasured candidates, if we assume that the active learning screen has identified all the top-performing molecules, a simple statistical analysis using the same 720-compound batch size reveals that a random screen stands a 25% chance of sampling the top performing candidate identified by our four-round active learning screen after 48 rounds and a 50% chance after 97 rounds. While chemical intuition and prior experience can assist and accelerate the search process, they also have the potential to create bias and preconceptions that may cause promising but non-intuitive candidates to be overlooked.

### 4.2 Analysis of the top performing candidates identified in the active learning screen

We now proceed to conduct a deeper analysis of the top-performing candidates identified by the active learning screen. First, we filtered 2880 experimentally assayed candidates for cytostatic or cytotoxic behavior using our confluency mask scores, as described in Section 2.2. This resulted in the removal of 303/2880 (10.5%) compounds from our screen, leaving a majority 2577/2880 (89.5%) as viable candidates. All following analyses will be conducted over viable candidates only, unless otherwise stated. The viability filter used to determine the viability of immunomodulator candidates for this work as described in Section 2.2 is more stringent than the previous work,^24^ and we will only compare the viable candidates in both works under these criteria.

Within this subset, we identified 167 novel immunomodulator candidates capable of mediating a 2-fold or more enhancement or suppression of at least one of the 12 objective functions, representing a 105% expansion in the palette of previously known immunomodulators with this level of NF-*κ*B/IRF modulation.^24^ We discovered molecules with unprecedented immunomodulatory capacity, including those capable of suppressing NF-*κ*B activity by up to 15-fold, elevating NF-*κ*B activity by up to 5-fold, and elevating IRF activity by up to 6-fold. Moreover, nine immunomodulators were observed to downregulate NF-*κ*B stimulation by more than 10-fold (i.e., fold change lower than 0.1), which also represents an unprecedented level of inhibition.^24^

Specifically, our top-performing enhancers of the NF-*κ*B response achieved fold improvements relative to agonist alone of 2.6-fold (LPS specialist), 5.5-fold (MPLA specialist), 2.8-fold (CpG specialist) and 2.9-fold (LPS, MPLA, CpG generalist). Our top-performing suppressors of the NF-*κ*B response achieved fold improvements relative to agonist alone of 0.1-fold (LPS specialist), 0.23-fold (MPLA specialist), 0.06-fold (CpG specialist), and 0.15-fold (LPS, MPLA, CpG generalist). Our top-performing enhancers of the IRF response achieved fold improvements relative to agonist alone of 5.9-fold (LPS specialist), 6.0-fold (MPLA specialist), 3.2-fold (cGAMP specialist), and 3.6-fold (LPS, MPLA, cGAMP generalist). The top-performing specialist candidates elevated NF-*κ*B activity by up to 110%, enhanced IRF activity by up to 83%, and suppressed NF-*κ*B activity up to 128% in comparison to the best-performing viable molecules identified in previous screening efforts over a more modest and (putatively) more immunologically-relevant 2674-compound library. ^24^ We also identified more generalists as compared to previous screening efforts. We discovered 8 NF-*κ*B generalists capable of inducing at least 1.5-fold enhancements in this pathway, 33 NF-*κ*B generalists capable of inducing at least 0.2-fold suppression of this pathway, and 15 IRF generalists capable of inducing at least 1.25-fold enhancements in this pathway. This represents a 36%, 236%, and 500% expansion in the number of known generalists from previous screening efforts.^24^

We present in Figure 4A the top two molecules for each of the 12 functional objectives identified by our screen, where we show the chemical structures and experimentally measured immunomodulatory profiles for each molecule. PME-4119 was identified as a top-performing NF-*κ*B suppressor generalist, as well as a top-performing MPLA NF-*κ*B suppressor specialist and a CpG NF-*κ*B suppressor specialist, showing that some potent generalists can also function as potent specialists. PME-5246 does not strongly enhance NF-*κ*B stimulation of any particular agonist, but it enhances NF-*κ*B stimulation with every agonist, meaning that it is a good generalist. However, potent IRF enhancer specialists appear to be poorer generalists due to their stronger specificities. Data on all 2880 molecules is presented in a CSV file in the Supporting Information that also provides the molecular SMILES strings and vendor from which the molecule was sourced.

**Figure 4:**
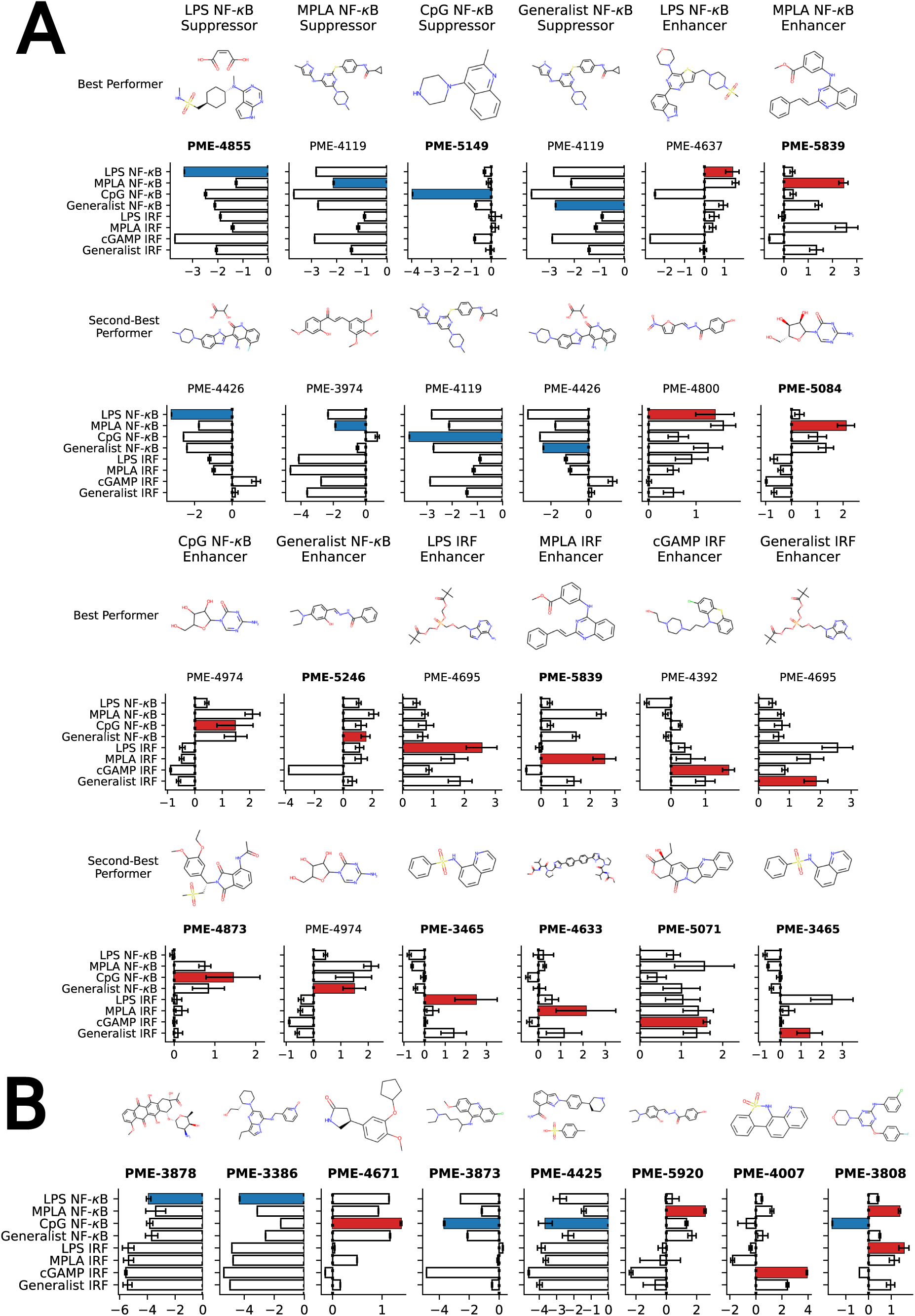
Top-performing immunomodulator candidates. (A) The two top-performing immunomodulator candidates in each of the 12 functional objectives. We present for each molecule its chemical structure along with their code names. A bar chart shows the experimentally measured log2-fold change in the immunomodulatory profile along all eight agonist-pathway combinations of interest. We highlight the immunomodulatory property that makes the candidate highly ranked in terms of activity enhancement (red) or suppression (blue). A full accounting of the measured performance of all 2880 molecules is presented in a CSV file in the Supporting Information. The 17 candidates with names highlighted in **bold text** were selected for additional characterization of their cytokine release profiles (note that PME-3465 and PME-5839 each appear top-ranked twice). (B) An additional eight candidates with outstanding immunomodulatory profiles that were selected for additional cytokine characterization. Although four of these molecules were determined to be non-viable in our confluency mask test for cytostatic or cytotoxic behavior (PME-3878, PME-3386, PME-5920, PME-4007), their exceptional immunomodulation capacity induced us to subject them to additional characterization.

We next traced the source of each top-performing molecule with a *>*2-fold enhancement/inhibition to identify that a significant number of these molecules come from the Microsource Spectrum Collection, Prestwick Chemical Library and Selleckchem FDA-approved Drug Library (Table S5). The common feature these three libraries share is that they all have a large portion of molecules that have been approved by some regulation institute, such as U.S. Food and Drug Administration (FDA), to be used as drugs or therapeutics. This shows that there is abundant repurposing potential for approved drugs to be used as immunomodulators.

We also computed the Tanimoto molecular similarity between each high-profile molecule with a *>*2-fold enhancement/suppression to identify the most similar molecule within the 2674 molecule data from our previous screen that was used to train the initial GPR model.^24^ The Tanimoto similarity metric quantifies the proportion of chemical substructures shared in a pair of molecules as a value between 0 and 1. The higher the Tanimoto similarity, the more substructures are shared between the molecules. A histogram of the Tanimoto similarity scores between the top performers and the most similar initial training molecule demonstrates significant support at low similarity values indicating that the active learning search has moved into new regions of space and is not simply sampling in the close vicinity of the training data (Figure S9).

Taken together, these results demonstrate the value of the active learning screen over large candidate libraries in efficiently identifying large numbers of novel small molecule candidates with high immunomodulatory activity.

### 4.3 Data-driven inference of chemical design rules

The active learning screen furnished immunomodulation measurements for 2880 new candidate molecules. Combining these with the 2674 compounds screened in previous study,^24^ we possess a rich data set of labeled immunomodulatory activity for 3560 compounds after removing non-viable and duplicated compounds. We then sought to interrogate these data to extract interpretable design rules for the immunomodulatory activity based on the molecular structure. It can be challenging to extract interpretable understanding of structure-function relationship learned by the GPR surrogate model. To furnish more comprehensible structure-function relations, we employed LASSO regression to train an interpretable linear model regressing the log2-fold change in immunomodulatory activity conditioned upon the presence or absence of particular chemical fragments or functional groups. In doing so, we exchange nonlinearity, complexity, and accuracy of the GPR for interpretability in the LASSO model predictions. The LASSO regularization term promotes sparse regression models in which many of the learned regression coefficients shrink to precisely zero (*θ_k_* = 0), and the remaining non-zero coefficients can be interpreted as pertaining to those chemical features that are the strongest determinants of observed immunomodulatory behaviors. The simple structure of the model means that the sign of the learned non-zero weights indicates the direction of immune response regulation. Specifically, chemical groups with a value of *θ_k_ >* 0 are associated with enhancing modulation and can be regarded as enhancer promoters. Conversely, chemical groups with a value of *θ_k_ <* 0 are associated with suppressing modulation and can be regarded as suppressor promoters.

In Figure 5 we present in non-ascending order of magnitude, the up to six non-zero regression coefficients for LASSO models fitted to each of the eight agonist-pathway combinations of interest. These weights can be interpreted as being associated with the features that have the highest predictive power for the log2-fold change in immunomodulatory activity in each of the eight agonist-pathway combinations. The chemical fragments pertaining to each regression coefficient are denoted by codes starting with “fr_” and followed by letters denoting the chemical groups they are quantifying. A glossary of all chemical fragment codes is provided as a CSV file within the Supporting Information and Figure S5 supplies a full accounting of all non-zero coefficients for all eight LASSO regression models.

**Figure 5:**
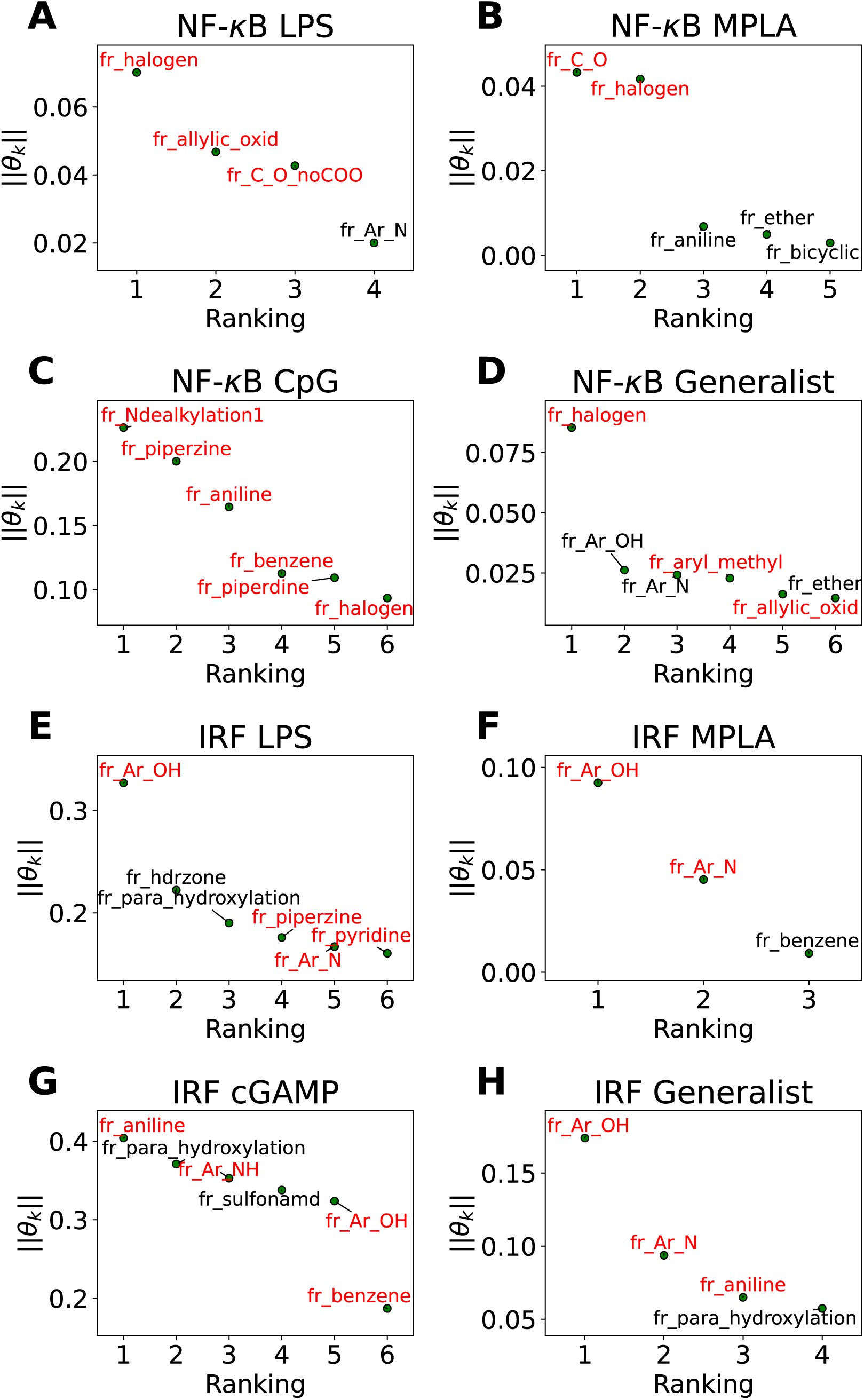
Immunomodulator design rules for each of the eight agonist-pathway combinations exposed by LASSO regression. Illustration of the up to six largest magnitude non-zero regression coefficients for LASSO linear regression models to predict the log2-fold change in immunomodulatory activity as a function of the presence or absence of particular molecular fragments or functional groups. The features with positive weights are displayed in black text, while the ones with negative weights are displayed in red text. Positive weights imply that there is a positive correlation between the feature values and enhanced immunomodulation, while negative weights indicate a positive correlation between the feature values and inhibitory immunomodulation. The chemical fragments pertaining to each regression coefficient are denoted by codes and a full glossary is available in the Supporting Information.

Our analysis reveals a number of interesting design rules. First, the halogen moiety “fr_halogen” appears as a negative, top-ranked fragment in all four of the NF-*κ*B specialist and generalist LASSO models. In particular, in the LPS and MPLA specialist and NF-*κ*B generalist categories it ranked among the top two. This indicates that the presence of halogen groups in immunomodulators is predictive of suppression of the activity of this pathway, especially immune responses activated by TLR4 agonists such as LPS and MPLA. This finding is aligned with several top performing candidates, some of which are shown in Figure 4A: a top-performing NF-*κ*B suppressor generalist and LPS specialist, PME-4426, has a fluoride group and PME-3873 and PME-4392, which are both NF-*κ*B suppressors, have chloride groups. Second, aromatic heteroatom moieties, including aromatic nitrogen “fr_Ar_N”, aromatic amine “fr_Ar_NH”, and aromatic hydroxyl group “fr_Ar_OH” appear frequently in multiple LASSO models with at least one of them retained in seven out of eight LASSO models, with the MPLA NF-*κ*B specialist being the only exception. Interestingly, the weights of these aromatic heteroatom moieties are positive for all NF-*κ*B LASSO models and negative for all IRF LASSO models. This result indicates that the presence of aromatic nitrogen/amine/hydroxyl group is predictive of enhancement of the immune activation in NF-*κ*B pathway, while it is also predictive of suppression of the activity of IRF pathway. Since the suppression of IRF pathway is of limited clinical significance, the information suggests elimination of these aromatic heteroatom moieties in IRF enhancer immunomodulators to optimize their potency. For example, PME-5246 and PME-4800, two NF-*κ*B suppressors, both possess aromatic hydroxyl groups, and PME-4974, which possesses an aromatic amine group, enhance NF-*κ*B responses while suppressing IRF responses in general. Third, the carbonyl moiety “fr_C_O_noCOO” appears as the third most influential fragment in the LPS NF-*κ*B specialist LASSO model. The sum of carbonyl and carboxyl moiety “fr_C_O” appears as a top-ranked fragments in the MPLA NF-*κ*B specialist and CpG NF-*κ*B specialist models with negative and positive weights, respectively. (Chemical fragments not ranked among top six illustrated in Figure 5 are presented in Figure S5). This indicates that the presence of carbonyl and carboxyl moieties appears to be predictive of suppression of the immune activity in NF-*κ*B pathway stimulated by TLR4 agonists such as LPS and MPLA, while it is predictive of enhancement of immune responses for CpG NF-*κ*B specialists. As a representative example, PME-4873, having three carbonyl groups in the structure, can greatly enhance NF-*κ*B response with CpG while it is a weak suppressor for NF-*κ*B response with LPS.

Overall, this analysis provides insights into understanding the characteristics of a molecule that promote specific immunomodulatory behaviors. These design rules are of value in advancing understanding of the possible modes of action of these molecules, suggesting how one might modify the structure of a particular immunomodulator to boost its performance in a particular immunomodulation goal, and in guiding how one might augment future candidate libraries to enrich them in molecules predicted to have promising immunomodulatory behaviors.

### 4.4 Validating top-performing immunomodulators with cytokine release profile measurements

Our high-throughput active learning screen already demonstrated the capacity of these immunomodulators to enhance or inhibit NF-*κ*B and IRF responses. However, these transcriptional activity measurements only provide an overall representation of the immunomodulatory behavior. In order to obtain a more detailed understanding of the immune signaling behaviors, we subjected 17 of the top-performing immunomodulator candidates identified in our active learning to a low-throughput assay measuring cytokine release profiles within primary cells (Figure 4). Specifically, we measured modulator’s ability to change cytokine secretion of murine bone marrow derived dendritic cells (BMDCs) stimulated with LPS, CpG and cGAMP. For this validation assay, MPLA was not tested as both LPS and MPLA are TLR4 agonists and one TLR4 agonist was determined to be sufficient for preliminary validation. The 17 top candidates were primarily selected from the top-performing immunomodulators in each of the 12 objectives – PME-5071, PME-4855, PME-4671, PME-4633, PME-4873, PME-3873, PME-5149, PME-4425, PME-3465, PME-5246, PME-5839, PME-3808 and PME-5084 – plus four molecules that were determined to be non-viable in our confluency mask test for cytostatic or cytotoxic behavior, but exhibit an exceptional immunomodulatory profiles – PME-3878, PME-3386, PME-5920 and PME-4007 (Figure 4B). Among these four additional candidates, two demonstrated potent inhibition – PME-3878 and PME-3386 – and two are potent enhancers – PME-5920 and PME-4007. The main results of this secondary screen are presented in Figure 6. Full data, including negative controls demonstrating that immunomodulators do not modulate cytokine secretion profiles in the absence of agonist, are presented in Figures S7, S8, and S12.

**Figure 6:**
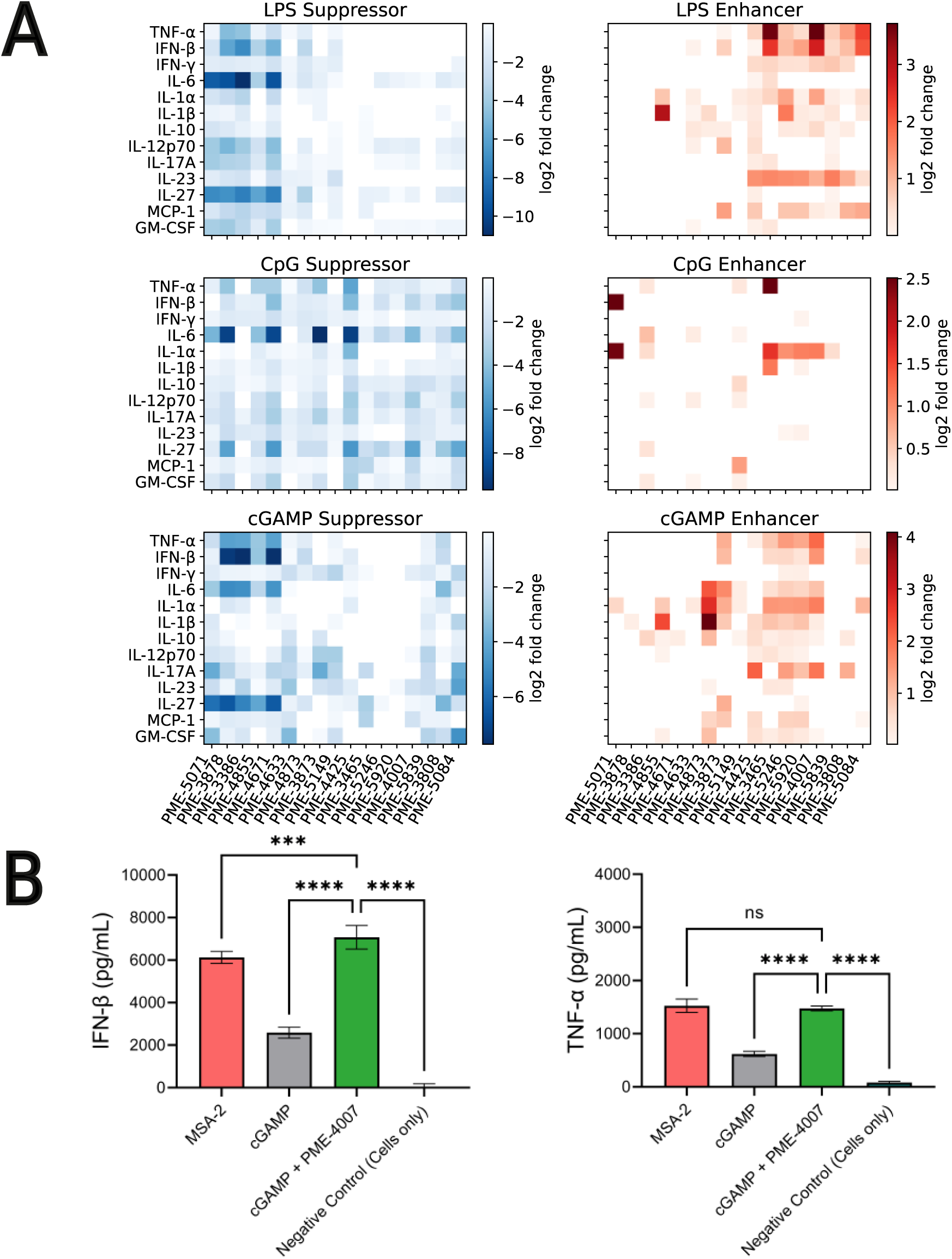
Low-throughput measurement of cytokine release profiles within primary cells of 17 top-performing candidates. (A) Immunomodulation of 17 selected top-performing candidates over the release profiles of 13 cytokines activated by LPS, CpG and cGAMP, shown as suppression and enhancement, repectively. The extent of immunomodulation is visualized as color-maps depicting the log2-fold change values, in form of heat-maps contrasting corresponding cytokines and immunomodulator indices. We are primarily interested in TNF-*α* and IFN-*β*, since NF-*κ*B activation is correlated with increases in proinflammatory cytokines such as TNF-*α*, whereas IRF activation is related to production of IFN-*β*. The top ranked immunomodulators show substantial capabilities in enhancing and inhibiting the production of TNF-*α* and IFN-*β*, as well as other cytokines such as IL-6, IL-27 and IL-1*α*. (B) Co-delivery of PME-4007 with cGAMP increases IFN-*β* secretion by more than three-fold relative to cGAMP alone. MSA-2 is a potent STING agonist that can stimulate IFN-*β* more strongly than cGAMP. While with the modulation induced by PME-4007 at a low concentration (2 *µ*M), the stimulation of cGAMP (10 *µ*g/mL) is enhanced to be significantly stronger than that induced by MSA-2 (10 *µ*g/mL). PME-4007 also enhances TNF-*α* secretion and helps reach a comparable level of that induced by MSA-2. Statistical analyses between agonist and modulator versus agonist alone were performed by a one-way ANOVA test (\**p* < 0.05, \*\**p* < 0.01, \*\*\**p* < 0.001, \*\*\*\**p* < 0.0001).

Immunologically, NF-*κ*B activation is correlated with increases in proinflammatory cytokines such as TNF-*α*, whereas IRF activation is related to production of IFN-*β*.^6–8,43^ Based on the results of our previous screen,^24^ we hypothesized that the enhancement or suppression of transcriptional activity induced by the immunomodulators should be associated with the increase or decrease in the production of relevant cytokines. Thus, we focused on the immunomodulation of the release of TNF-*α* and IFN-*β*.

We observed from our secondary screening results that six immunomodulators – PME-3878, PME-3386, PME-4671, PME-4425, PME-3465, and PME-4007 – out of the 17 top-ranked candidates demonstrated significant capacity to modulate the secretion of TNF-*α* and IFN-*β* when co-delivered with LPS, CpG, or cGAMP (Figure 6A). Taken together, this cytokine validation assay demonstrated that the immunomodulators identified by our active learning-guided pipeline upregulate TNF-*α* production by over 10-fold, downregulate TNF-*α* production by over 16-fold, and upregulate IFN-*β* production by over 6-fold.

Three immunomodulators stand out as of particular interest. PME-3878 and PME-3386 are two candidates discovered in our active learning screen as top-performing generalist inhibitors of NF-*κ*B as well as top-performing specialist inhibitors for NF-*κ*B when treated with LPS. PME-4007 is a candidate that is a top-performing specialist enhancer of the IRF when treated with cGAMP. PME-3878 and PME-3386 inhibit TNF-*α*, IL-6, and IFN-*β* production for nearly all agonists considered (Figure 6A). Suppressing immunomodulators like these can be used as potential adjuvants for prophylactic vaccines or therapeutics that benefit from minimizing pro-inflammatory cytokines. In contrast, PME-4007 is a moderate to strong enhancer of the TNF-*α*, IFN-*β*, IL-1*α*, and/or IL-17A responses in the presence of LPS, CpG, or cGAMP (Figure 6A). cGAMP is a pattern recognition receptor agonist that acts through the STING pathway.^15^ Immunomodulators that enhance IFN-*β* production through the STING pathway are of particular interest in promoting antiviral defense and anti-tumor immunity thorough T cell cross priming. ^62^

We further subjected our leading IFN-*β* inducing compound, PME-4007, to additional comparisons of its cytokine profile in the presence of cGAMP to MSA-2, a recently identified STING agonist.^63^ MSA-2 was discovered via a high throughput process involving over two million compounds and is more potent than cGAMP. As illustrated in Figure 6B, we observed MSA-2 to induce an IFN-*β* secretion that is significantly higher than that induced by cGAMP at the same concentration (10 *µ*g/mL). Furthermore, when PME-4007 was added in a low concentration (2 *µ*M) in combination with cGAMP, it increased IFN-*β* levels by approximately 3-fold relative to cGAMP treated cells and IFN-*β* was statistically significantly elevated relative MSA-2 treated cells. PME-4007 also enhanced TNF-*α* secretion stimulated by cGAMP to a level that is comparable to that stimulated by MSA-2. Our *in vitro* screen identified an immunomodulator that can be combined with a commonly used, naturally occurring STING agonist to induce similar immunological profiles to a best-in-class STING agonist.

## 5 Conclusions

Co-delivery of immunomodulators with PRR agonists presents a powerful means to reduce inflammation or otherwise modulate innate immune stimulation by enhancing or suppressing innate immune signaling pathways, and offers a route to improving vaccines by reducing adverse side-effects and cancer therapies by enhancing the magnitude of the immune response. Small molecules present attractive immunomodulator candidates with high synthetic accessibility and reduced immunogenic potential compared to biologics. The vast size of the drug-like small molecule design space makes strategies to maximize the utility of each experimental assay extremely valuable in rationally and effectively traversing this space. In this work, we combined we constructed a data-driven QSAR model combining deep representational learning, Gaussian process regression, and Bayesian optimization to guide high throughput experimental screening of a library of 139,998 commercially available candidate small molecules. After conducting four rounds of an active learning search that screened 2880 molecules (*∼*2% of the search space) we identified novel immunomodulator candidates capable of suppressing NF-*κ*B activity by up to 15-fold, elevating NF-*κ*B activity by up to 5-fold, and elevating IRF activity by up to 6-fold. The top-performing candidates furnished a 110% improvement in NF-*κ*B activity, 83% improvement in elevating IRF activity, and 128% improvement in suppressing NF-*κ*B activity, and we also identified 167 novel immunomodulators with at least 2-fold enhancement or suppression over transcription factor activity of interest – representing a 105% increase in the total number of known immunomodulators with this level of activity – and nine novel immunomodulators with at least 10-fold activity modulation, while this level of activity modulation is not previously observed with the immunomodulator candidates in the previous work. ^24^ Additional characterization of the cytokine release profiles of the top 17 candidates demonstrated their ability to substantially modulate key cytokines such as TNF-*α*, IL-6, and IFN-*β*, in combination with particular PRR agonists. One particular candidate, PME-4007, was observed to produce a 3-fold increase in IFN-*β* production when co-delivered with the STING agonist cGAMP, which is comparable with the recently developed STING agonist MSA-2 identified via a large two-million-compound high throughput screen. ^63^ Finally, we performed a *post hoc* analysis of the high-throughput screening data using interpretable linear models to extract interpretable design rules linking the presence or absence of particular functional groups to immunomodulatory performance. Of particular interest, we found halogen moieties to be correlated with suppression of NF-*κ*B activity, and carbonyl and carboxyl moieties with suppression of NF-*κ*B activity in pathways activated by TLR4 agonists such as LPS and MPLA, but suppression of the IRF pathway.

In future work, we plan to conduct more detailed characterization of the top performing small molecule candidates identified in this screen including *in vivo* testing and kinetic measurements to unveil their mechanism of action. We also plan to expand our screen to larger candidate libraries, including those enriched in molecules adhering to the design rules extracted from our analysis. Finally, we also plan to improve our screen to incorporate additional constraints on physical properties of the immunomodulators such as water solubility and synthetic accessibility, to better facilitate their incorporation into vaccine formulations and delivery as vaccines and therapeutics.

## Supporting information

descriptors.csv

fragments-meaning-retained.csv

hts-2880-modulator.csv

lib-140K.csv

## Acknowledgement

This work was supported by National Institute of Allergy and Infectious Diseases of the NIH under the Discovery of Adjuvant Program: award NIH 75N93019C00041. The content is solely the responsibility of the authors and does not necessarily represent the official views of the NIH. This work was completed in part with resources provided by the University of Chicago Research Computing Center. We gratefully acknowledge computing time on the University of Chicago high-performance GPU-based cyberinfrastructure supported by the National Science Foundation under Grant No. DMR-1828629.

## Data And Code Availability Statement

We present a full accounting of the integrated 140K-compound library used to train the VAE model (with source library information and SMILES); the screened 2880-compound library (with SMILES, SELFIES, modulation fold change and viability score); the glossary of chemical fragments code names corresponding to their chemical interpretations; Python implementation of the VAE model; Python implementation of the Bayesian optimization model and the multi-objective Kriging believer sampling strategy; Python implementation of visualization code used for plotting data. The code repository is available at https://github.com/oytang/Supporting-data-for-Data-driven-discovery-of-innate-immunomodulators-Supporting

## Supporting Information Available

The following files are available as Supporting Information free of charge.

- si.pdf: Consolidated Supporting Information.
- lib-140K.csv: The integrated 140K-compound library used to train the VAE model (with source library information and SMILES).
- hts-2880-modulator.csv: The screened 2880-compound library (with SMILES, SELFIES, modulation fold change and viability score).
- fragments-meaning-retained.csv: The glossary of chemical fragments code names corresponding to their chemical interpretations, with irrelevant fragments removed.
- descriptor.csv: A spreadsheet listing all 85 descriptors, within which we designate them as “Irrelevant” or “Redundant” if they were eliminated in the corresponding feature selection step, and “Effective” if they were retained in further investigation.

## TOC Graphic and Synopsis

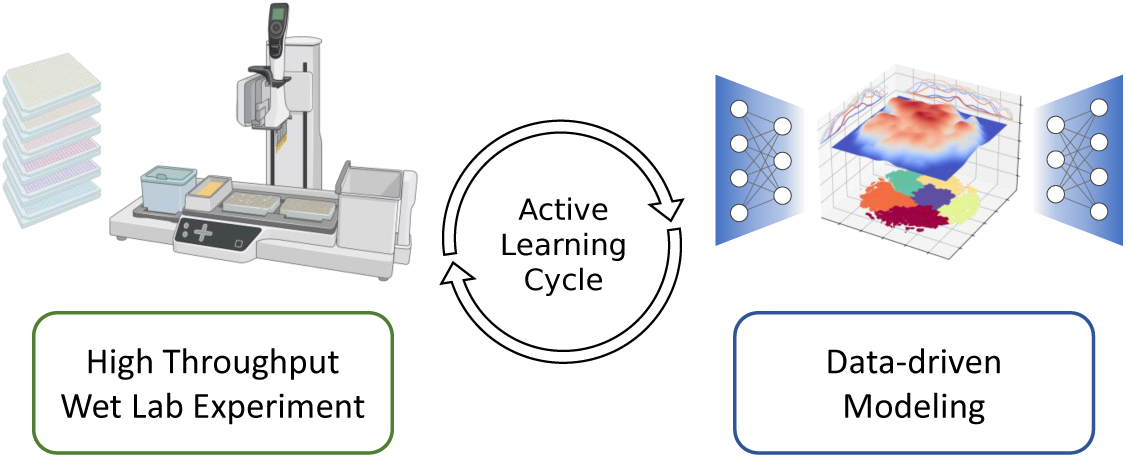

We combine high-throughput wet lab experimentation and data-driven computation in a closely coupled active learning loop in order to identify novel molecules with exceptional properties.

