## Supplementary Information for "Data-driven discovery of innate immunomodulators via machine learning-guided high throughput screening"

**S3 Supplementary Figures** **26**

**S4 Supplementary Tables** **30**

**References** **31**

### S1 Supplementary Computational Methods

#### S1.1 Definition of ∼1M compound augmented ZINC library

We defined an augmentation of the ZINC small molecule library[1–4] to train our VAE network for deep representational embeddings of our immunomodulator candidates. We combined the 2,674 molecules employed in our prior screening work,[5] with a subset of 924,870 molecules from the ZINC libraries[1–4] comprising those molecules that containing a biphenyl scaffold, inspired by the structure of previous small molecule immunomodulator discovery of Honokiol.[6] We also incorporated some other generic molecular libraries from various vendors. As a result, our augmented ZINC library consists of 1,262,866 small molecule compounds in total. A list of the molecular libraries compiled to define the initial augmented ZINC library is provided in Table S1.

We then filtered the library under a number of criteria. First, we represented each compound as simplified molecular-input line-entry system (SMILES) strings.[7–10] We then canonicalized these SMILES representations and removed duplicates. Second, we translated each SMILES string into self-referencing embedded strings (SELFIES) as a more robust representation of molecules that are guaranteed to define valid chemical structures.[11] We then eliminated compounds that produced an inconsistent SMILES string upon back-translation from the SELFIES representation to ensure a strict one-to-one mapping between SMILES representations and SELFIES representations of the compounds, thus enforcing a strict one-to-one mapping between SELFIES representations and the structure of the compounds. With the first two steps of filtering, we removed 8734 entries and were left with 1,254,132 compounds in the library. Third, we capped the maximum SELFIES string length to 137 characters and eliminated compounds containing tokens that appear in fewer than 500 compounds. Since the SELFIES strings will ultimately be represented to the VAE input layer as fixed-length vectors, this step constrained the dimension of the representation space so as to limit the size of the network and its associated training costs. These constraints resulted in the elimination of only 5468 compounds, corresponding to fewer than 0.5% of candidates in the immunomodulator library (Section S1.2). After performing these filtering operations, we were left with 1,248,664 compounds in the augmented ZINC library.

Finally, we digitized the SELFIES representations into one-hot matrices of dimension 137-by-55, where 137 represents the maximum possible string length and 55 are the number of possible SELFIES tokens, including the null token. SELFIES strings less than the maximum length are padded with null tokens. We then flattened the one-hot matrices into 7535-element vectors to provide a fixed length representation of each molecule to be passed to the VAE.

Table S1: **Compound libraries used to assemble the augmented ZINC library.**

| Company/Institution | Compound Library | # Compounds | URL |
|---|---|---|---|
| UChicago | Our Prior Screen Work[5] | 2674 | N/A |
| Analyticon | Synthetic Screening Compounds | 5000 | URL |
| ChemBridge | DS550-3 | 29882 | URL |
| ChemBridge | ES550-1 | 49966 | URL |
| ChemBridge | ES550-2 | 49958 | URL |
| ChemRoutes | ChemArray | 21135 | URL |
| Life Chemicals | Pre-plated Diversity Sets PS4 | 50200 | URL |
| Life Chemicals | Pre-plated Diversity Sets PS5 | 50168 | URL |
| Life Chemicals | Biologically Active Compound Library | 7991 | URL |
| MedChemExpress | STAT Inhibitors | 58 | URL |
| Micro Source | The Spectrum Collection | 1975 | URL |
| Prestwick Chemical | Prestwick Chemical Library | 1276 | URL |
| Selleckchem | FDA-approved Drug Library | 2667 | URL |
| Selleckchem | STAT Inhibitors | 18 | URL |
| UCSF SMDC | Various Compound Libraries | 65790 | URL |
| **ZINC[1–4]** | Biphenyl Scaffold | 924870 | URL |
| | **Sum** | **1262866** | |
| | **Sum (After Filtering)** | **1248664** | |

#### S1.2 Definition of ~140k compound immunomodulator candidate library

We selected seven generic commercial small molecule compound libraries to define an initial pool of 139,998 candidate immunomodulators to draw candidates from and bring to high throughput screening experimentation. These libraries were a subset of our previously assembled 1,248,664 compound augmented ZINC library. These screening compound libraries were designed with the intention of enabling cell-based and target-based high throughput screening initiatives by making a diverse range of small molecules readily available, which make it easier for us to access the molecules for screening experiments. This immunomodulator candidate library went through the same filtering steps as introduced in Section S1.1, with the only exception that we were not using a new set of SELFIES tokens, but we used the token dictionary we used for the augmented ZINC library. This is to ensure the mapping between the network parameters of VAE and the one-hot encodings of compounds. We transformed the SELFIES representations into one-hot matrices, with a dimension of 137-by-55 then flattened the one-hot matrices into 7535-element vectors to obtain a standardized representation of each molecule. A list of the molecular libraries compiled to define the initial immunomodulator candidate library is provided in Table S2. A CSV file providing SMILES strings of all compounds of the 139,998 compound library is provided in the Supporting Information.

Table S2: **Compound libraries used to assemble the immunomodulator candidate library.**

| Company | Compound Library | Number of Molecules | URL |
|---|---|---|---|
| ChemBridge | DS550-3 | 29547 | URL |
| ChemBridge | ES550-3 | 49584 | URL |
| Life Chemicals | Biologically Active Compound Library | 7938 | URL |
| Life Chemicals | Pre-plated Diversity Sets PS4 | 50233 | URL |
| Micro Source | The Spectrum Collection | 1821 | URL |
| Prestwick Chemical | Prestwick Chemical Library | 1233 | URL |
| Selleckchem | FDA-approved Drug Library | 1539 | URL |
| | **Sum (After Filtering)** | **139998** | |

#### S1.3 Deep representational learning using variational autoencoders (VAEs)

##### S1.3.1 Model architecture

We employ a VAE architecture inspired by prior work by Aspuru-Guzik and co-workers[11,12] and implemented in PyTorch.[13] The encoder consists of three fully-connected (FC) layers that passes into a fourth fully-connected bottleneck layer corresponding to the latent space embedding. For the decoder, we employed a stack of gated recurrent units (GRU).[14] The architecture of the VAE is illustrated in Figure S1. The hyperparameters of the model architecture were optimized over the ranges reported in Table S3.

Table S3: **Architecture and hyperparameter optimization ranges for the VAE.**

| Hyper-parameter description | Value | Optimization range |
|---|---|---|
| The number of neurons in the first FC layer | 500 | [250, 1000] |
| The number of neurons in the second FC layer | 200 | [200, 500] |
| The number of neurons in the third FC layer | 100 | [50, 250] |
| Latent space dimension | 100 | [3, 250] |
| The number of recurrent layers of GRU | 3 | [1, 3] |
| The number of features in the hidden state of GRU | 200 | [20, 400] |

##### S1.3.2 Model training over augmented ZINC library

The VAE loss function $\mathcal{L}_{\text{VAE}}$ comprises two components: (1) the reconstruction loss $\mathcal{L}_{\text{Rec}}$ measured by the cross-entropy between the input and output one-hot SELFIES vectors to enforce reconstruction fidelity and (2) the Kullback-Leibler divergence (KLD)[15] $\mathcal{L}_{\text{KLD}}$ of the latent vectors relative to the standard normal distribution to regularize the latent space,[12,16]

$$\mathcal{L}_{\text{VAE}} = \mathcal{L}_{\text{Rec}} + \mathcal{L}_{\text{KLD}}, \tag{S1}$$

where,

$$\mathcal{L}_{\text{Rec}} = -\sum_{i=1}^{N}\sum_{j=1}^{D} x_{ij}\log\left(\hat{x}_{ij}\right) + \left(1 - x_{ij}\right)\log\left(1 - \hat{x}_{ij}\right), \tag{S2}$$

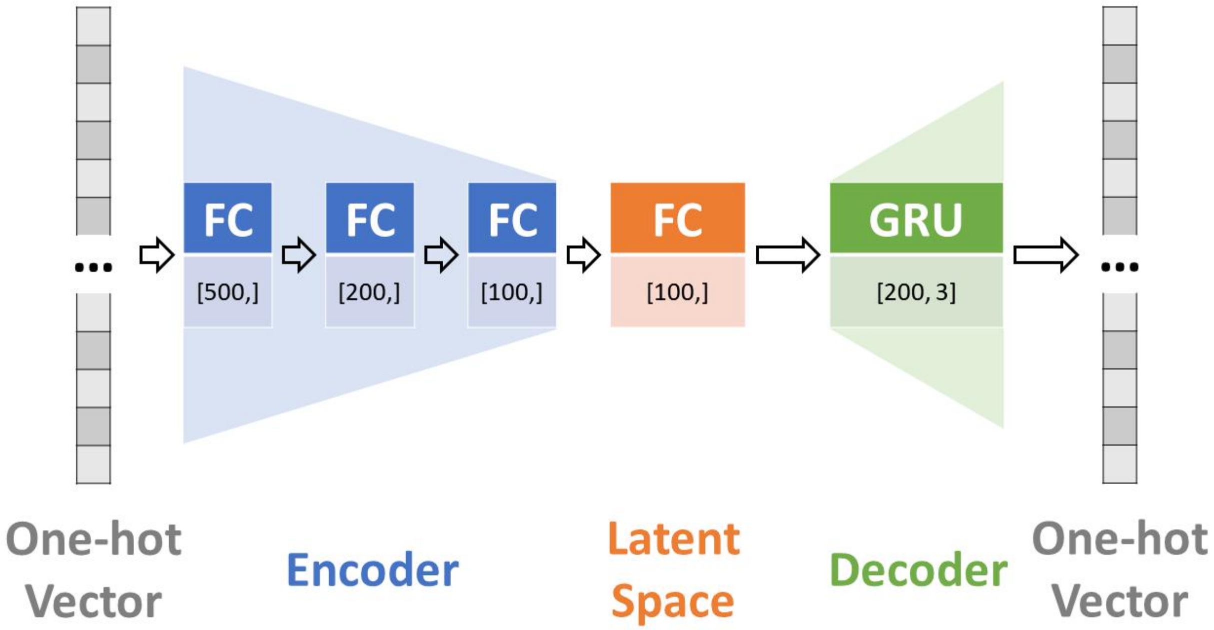

Figure S1: **Architecture of VAE for deep representational learning of molecular latent space.** The input and output of the VAE model are 7535-element one-hot vector that are one-to-one with the SELFIES string representation of each molecule. The encoder stack comprises three fully-connected feedforward layers with a 500-200-100 architecture. A bottleneck layer comprising 100 neurons defines the 100D latent space embedding. The decoder comprises three stacked GRUs each containing 200 neurons.

where $N$ is the number of samples, $D$ is the dimension of the data, $x_{ij}$ is the $j^{\text{th}}$ component of the $i^{\text{th}}$ input vector, and $\hat{x}_{ij}$ is the corresponding reconstructed output of the autoencoder, and,

$$\mathcal{L}_{\text{KLD}} = -\frac{1}{2}\sum_{j=1}^{J}\left(1+\log(\sigma_j^2)-\mu_j^2-\sigma_j^2\right), \tag{S3}$$

where $\mu_j$ and $\sigma_j^2$ are the mean and variance of the $j^{\text{th}}$ element of the latent vector $z$, respectively, and $J$ is the dimensionality of $z$.

Network training is conducted by minimizing Equation S1 using the Adam optimizer.[17] We first train the VAE over the augmented ZINC library for 10,000 epochs. To guard against posterior collapse,[18] we employ an initial KLD loss coefficient of $\alpha = 10^{-2}$ that we schedule to reduce to $\alpha = 10^{-3}$ at epoch 2000, and to $\alpha = 10^{-4}$ at epoch 5500. The training hyperparameters comprising batch size and learning rates were optimized over the ranges reported in Table S4. We assess model performance by computing the exact reconstruction accuracy defined as the fraction of molecules in the training set whose SELFIES strings can be reconstructed with 100% fidelity. Since the intended application of the trained VAE is to define a latent space embedding of our candidate molecules to support an active learning search, we use the decoder in service of learning this latent embedding but, in the present work, do not make use of its generative capacity to extend the model beyond the training data and produce novel synthetic molecules. As such, we assess model performance on the training data rather than the usual practice of employing a hold-out test set. Our model achieves an exact reconstruction accuracy of 97.3% at epoch 9032 after more than 1344 GPU-hours of training. This high accuracy indicates that the network has discovered a 100D latent space embedding that preserves the salient information necessary for accurate molecular reconstruction, and we terminate training at this point. The training curves for the model are illustrated in Figure S2.

Table S4: **VAE training hyperparameters and optimization ranges.**

| Training parameter | Value | Optimization range |
|---|---|---|
| Batch size | 2500 | [2000, 3000] |
| Initial learning rate of the encoder | 0.0001 | [0.0001, 0.01] |
| Initial learning rate of the decoder | 0.0001 | [0.0001, 0.01] |

#### S1.4 Gaussian process regression (GPR) surrogate models

In our present application, we seek to identify immunomodulators capable of enhancing or suppressing the immune activity of a certain pathway when the pathway is activated by a certain agonist. This leads to our optimization in maximizing or minimizing the immunomodulation values as fold changes. To guide our search in the chemical design space towards profitable regions, we train a surrogate model to predict the fitness of immunomodulator candidates that have not been experimentally tested. We include an uncertainty measure that reflects our confidence in the predictions, enabling us to select points in the

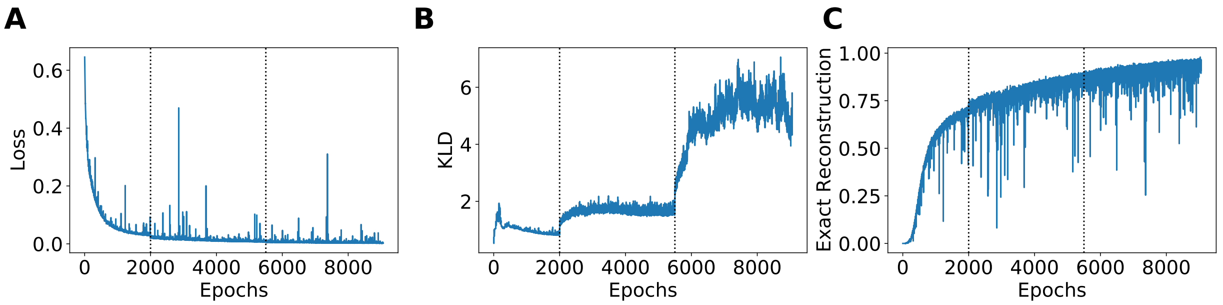

Figure S2: **VAE training curves over the augmented ZINC training library.** (a) VAE loss function (Equation S1). (b) KLD component of VAE loss (Equation S3). The dashed vertical lines in each panel denote the epochs at which the KLD loss coefficient $\alpha$ was tuned from an initial value of $10^{-2}$ to $10^{-3}$ then $10^{-4}$. (c) Exact reconstruction accuracy reporting the fraction of molecules in the training data whose SELFIES representations are exactly reconstructed by the trained network.

space that balance both high predicted fitness (exploitation) and high uncertainty (exploration). For this supervised regression task, we use Gaussian Process Regression (GPR) as our surrogate model, as it is a non-parametric Bayesian regression technique that provides built-in uncertainty estimates.[19] We have chosen to use the radial basis function (RBF) kernel (also known as the squared exponential kernel) as the covariance function within the GPR implementation in scikit-learn,[20]

$$k(x_i, x_j) = \sigma^2 \exp\left(-\frac{|x_i - x_j|^2}{2\ell^2}\right), \tag{S4}$$

where $x_i$ and $x_j$ are input data points, $\sigma^2$ is the signal variance, $\ell$ is the length scale of the kernel, and $|\cdot|$ represents the Euclidean distance between two points. To account for uncertainties associated with each experimental measurement $\sigma^n$, we adopt Tikhonov regularization[21] such that an uncertainty vector $\sigma = [\sigma_1, \sigma_2, \ldots, \sigma_n]^T$ is added to the diagonal of the kernel matrix $K$ during fitting. Although in each experiment, the standard deviation for each sample being tested was different, we took the average of the experimental errors of all samples $\overline{\sigma} = \frac{1}{n}\sum_{i=1}^{n}\sigma_i$ and fed into the GPR models as the uncertainty vector $\sigma = [\overline{\sigma}, \overline{\sigma}, \ldots, \overline{\sigma}]^T$.

As we were seeking immunomodulators with various capabilities – different kinds of immunomodulation (enhancement, suppression), different pathways (NF-$\kappa$B, IRF), and different agonists (LPS, MPLA, CpG, cGAMP) – we pursue a multi-objective optimization. As such, we constructed multiple GPR models, with each GPR corresponding to enhancement/suppression with a specific agonist on a specific pathway. In addition to specialists – immunomodulators that induce large changes in enhancement or suppression when co-delivered with one particular agonist – we also sought to identify generalists – immunomodulators that lead to large effects when co-delivered with any one of a group of agonists. We constructed GPR models for generalists using the arithmetic mean of the immunomodulation fold change values of corresponding specialist objectives. For example, as one of the objectives for enhancers of the NF-$\kappa$B response, the immunomodulation of generalist over LPS, MPLA and CpG agonists can be expressed as

$$M^{\text{NF-}\kappa\text{B Enhancer}}_{\text{generalist(LPS, MPLA, CpG)}} = \frac{1}{3}\left(M^{\text{NF-}\kappa\text{B Enhancer}}_{\text{LPS}} + M^{\text{NF-}\kappa\text{B Enhancer}}_{\text{MPLA}} + M^{\text{NF-}\kappa\text{B Enhancer}}_{\text{CpG}}\right), \quad \text{(S5)}$$

where $M^{\text{NF-}\kappa\text{B Enhancer}}_{\text{LPS}}$, $M^{\text{NF-}\kappa\text{B Enhancer}}_{\text{MPLA}}$ and $M^{\text{NF-}\kappa\text{B Enhancer}}_{\text{CpG}}$ are immunomodulation fold change values for modulator specialist with agonist LPS, MPLA and CpG, respectively. We initially have 14 objectives, each representing a GPR model: (1) Enhancers of the NF-$\kappa$B response (4 objectives): specialist for LPS agonist, specialist for MPLA agonist, specialist for CpG agonist, and generalist over LPS, MPLA, and CpG agonists; (2) Inhibitors of the NF-$\kappa$B response (4 objectives): specialist for LPS agonist, specialist for MPLA agonist, specialist for CpG agonist, and generalist over LPS, MPLA, and CpG agonists; (3) Enhancers of the IRF response (6 objectives): specialist for LPS agonist, specialist for MPLA agonist, specialist for CpG agonist, specialist for cGAMP agonist, generalist over LPS, MPLA, and CpG agonists, and generalist over LPS, MPLA, CpG and cGAMP agonists. We did not seek to identify inhibitors of the IRF response because of limited clinical significance of such immunomodulation. We did not optimize for enhancers or inhibitors of the NF-$\kappa$B response for specialist for cGAMP agonist because we knew from prior work that cGAMP does not strongly stimulate the NF-$\kappa$B pathway.[22] Following our screening experiment, we found that the specific CpG agonist we were using (CpG ODN 1826) demonstrated minimal IRF stimulation, as can be seen in Figure S6. As a result, we only report specialist for CpG agonist as enhancers and inhibitors of the NF-$\kappa$B response, and the 14 initial objectives were reduced to the 12 objectives as shown in Figure 1E.

#### S1.5 Multi-objective Bayesian optimization (BO)

After building GPR surrogate models to predict various forms of immunomodulation for every compound, our objective is to explore the design space by iteratively querying new compounds to experimentally test. This involves striking a balance between selecting the most promising candidates with high expected performance (exploitation) and potentially valuable candidates with high uncertainty associated with their prediction (exploration). To achieve this balance, we utilize Bayesian optimization (BO), which is a powerful black-box optimization technique that utilizes the posterior mean and uncertainties derived from GPR predictions to identify the most promising candidates for testing.[23] BO can optimize costly black-box functions by optimizing the surrogate objectives that are defined by GPR. An acquisition function is needed to determine the next best candidate for evaluation. To meet our goal of discovering more effective immunomodulators, we have chosen the Expected Improvement (EI) acquisition function acquisition function.[24] This function guides the selection of candidates that are predicted to result in maximum expected gains in fitness. The EI acquisition function can be expressed as,

$$\mathrm{EI}(\mathbf{x}) = \begin{cases} (\mu(\mathbf{x}) - f(\mathbf{x}^+) - \xi)\Phi(Z) + \sigma(\mathbf{x})\phi(Z), & \text{if } \sigma(\mathbf{x}) > 0, \\ 0, & \text{otherwise} \end{cases} \tag{S6}$$

where $\mathbf{x}$ is the input point, $\mu(\mathbf{x})$ and $\sigma(\mathbf{x})$ are the mean and standard deviation of the surrogate model prediction at $\mathbf{x}$, $f(\mathbf{x}^+)$ is the best function value observed so far, $\xi$ is a trade-off parameter that balances exploration and exploitation, $\Phi$ and $\phi$ are the standard normal cumulative distribution function and probability density function, and $Z = \frac{\mu(\mathbf{x})-f(\mathbf{x}^+)-\xi}{\sigma(\mathbf{x})}$ is the standard normal random variable. We select the next best candidate point $\mathbf{x}^*$ with the highest value of the acquisition function from the set of available candidates $\mathbf{x}_k$ that have not yet been sampled, i.e., $\mathbf{x}^* = \mathrm{argmax}_k \mathrm{EI}(\mathbf{x}_k; \mathbf{x}_k \notin \{\mathbf{x}_1, \mathbf{x}_2, \ldots, \mathbf{x}_n\})$. In this work, we employ $\xi = 0.01$ as a good general purpose choice commonly employed in the literature.[24,25]

Traditional multi-objective Bayesian optimization (MOBO) strategies typically aim to balance multiple objectives to efficiently map out Pareto optimal solutions.[26,27] In this work, this would be valuable in identifying potent generalists, but may sacrifice the discovery of potent specialists that reside at the "corners" of the Pareto frontier. Furthermore, some of our objectives are mutually incompatible, such as seeking to find enhancers and inhibitors for the same pathway with the same agonist. Accordingly, we adopted an alternative MOBO strategy in which we polled each of our 12 GPR models independently to collect the best candidates recommended by each model under the EI acquisition function and eliminated duplicates. To make efficient use of the experimental assay, we followed a batched sampling Kriging believer approach[28,29] wherein we asked each model for its next top-ranked candidates until we had collected a batch of 720 molecules for experimental testing. Under this protocol, we temporarily augment the training data with the molecules selected by the GPRs annotated with the GPR activity predictions, the GPR models are retrained on this augmented data, and the models polled again for their next top-ranked prediction under the EI acquisition function. This process is repeated until a batch of 720 molecules has been collated for experimental testing. This batched sampling approach sacrifices efficiency from an information theoretic perspective – each model is asked to choose a series of candidates without first receiving back experimental information on the previously selected candidates – but increases temporal efficiency by testing multiple molecules in an experimental batch.

Importantly, all selected molecules, regardless of their source GPR models, were subjected to comprehensive experimental testing to evaluate their immunomodulatory profiles across all agonist types and activation of both NF-$\kappa$B and IRF pathways. As such, once the experiments are complete, the activity entries in the augmented training data that were completed with the GPR activity predictions in service of the Kriging believer batching are corrected with the measured activities, and these data used to retrain all GPR model. In this manner, molecules selected by one GPR model are not only used to retrain that particular GPR model, but also used to retrain all other models.

#### S1.6 Inference of chemical design rules using LASSO regression

A comprehensive list of descriptors that RDKit is able to compute is available at `https://www.rdkit.org/docs/GettingStartedInPython.html#list-of-available-descriptors`, which includes a total of 208 2D descriptors. We elected not to include 3D descriptors since 3D molecular structures were not readily available for most of the compounds in our study. We selected those 85 descriptors denoting the occurrence of chemical groups for better interpretability. During feature selection, eight irrelevant descriptors were removed because they are invariant across all 3560 viable compounds in our study. To eliminate redundant descriptors, we looked at the 13 descriptors with Pearson correlation coefficients higher than 0.95 with any other descriptors. We then used clustering analysis to group highly correlated descriptors together into larger families or categories (Figure S3). In each group of highly correlated descriptors, we selected one descriptor as representative of the group. In this way, seven additional descriptors were marked redundant and removed. In sum, we retained 85-8-7 = 70 effective descriptors that were kept for further analysis. A CSV file listing all 85 descriptors is provided in the the Supporting Information, within which we designate them as "Irrelevant" or "Redundant" if they were eliminated in the corresponding feature selection step, and "Effective" if they were retained in further investigations.

Having defined the featurization for each immunomodulator, we construct a straightfor-

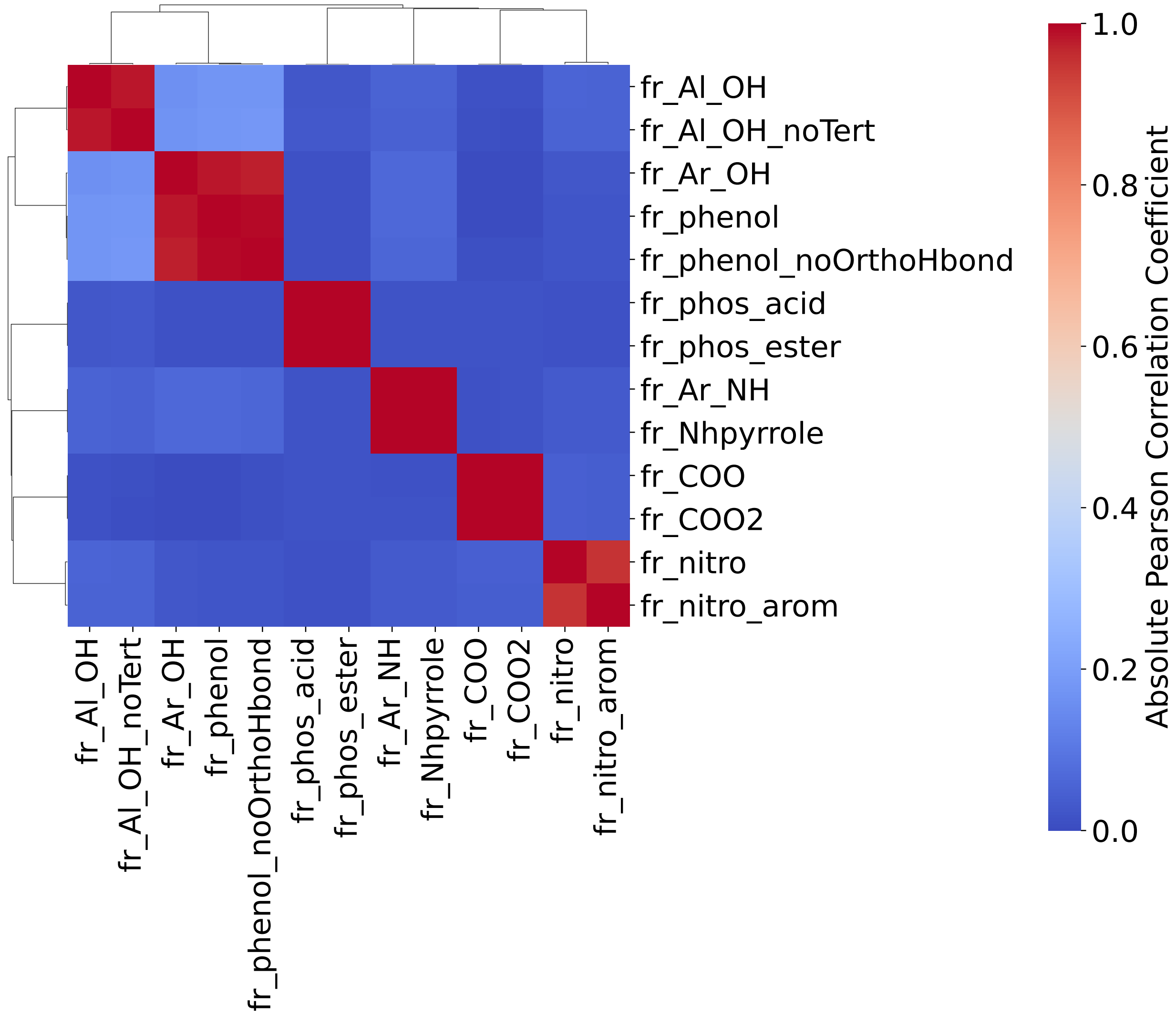

Figure S3: **Correlation heatmap and cluster dendrogram of highly correlated descriptors.** We selected 13 descriptors that had a Pearson correlation coefficient greater than 0.95 with at least one other descriptor. Clustering analyses were conducted to organize highly correlated descriptors into larger groups/families. In this way, we identified six descriptor groups with high correlation (red blocks shown in the clustered heat map). For each of the six descriptor groups, we select one representative descriptor. Thus, we retained six descriptors, namely "fr_Al_OH", "fr_Ar_OH", "'fr_phos_acid", "fr_Ar_NH", "fr_COO", "fr_nitro".

ward and interpretable model to predict the log2-fold change in immunomodulatory activity $\log_2 M$ as a linear function of the $k = 70$ standardized features $F_k$,

$$\log_2 M_{\text{predicted}} = \frac{1}{N}\sum_{N=1}^{3560} \log_2 M_{\text{exp}} + \sum_{k=1}^{70} \theta_k F_k, \tag{S7}$$

where $\frac{1}{N}\sum_{N=1}^{3560} \log_2 M_{\text{exp}}$ is the arithmetic mean of the log fold change in immunomodulatory activity over all $N = 3560$ experimental measurements and $\theta \in \mathbb{R}^{70}$ is a vector regression coefficients assigning weights to the different features. The coefficients $\theta$ are learned by minimizing the LASSO regression loss between the predicted and experimentally measured log2-fold change in immunomodulatory activities,

$$\mathcal{L}_{\text{LASSO}}(\theta; \alpha_L) = \frac{1}{2}\|\log_2 M_{\text{predicted}} - \log_2 M_{\text{exp}}\|_2^2 + \alpha_L \|\theta\|_1. \tag{S8}$$

The first term on the right side of the equation is the mean squared error between the predicted and experimental immunomodulation values, and the second term is the L1 regularization penalty term with hyperparameter $\alpha_L$. The $\|\cdot\|_2$ and $\|\cdot\|_1$ represent the L2 and L1 norms, respectively. The L1 regularization penalty term encourages the model to have fewer non-zero coefficients or parameters, thereby promoting a sparse model that can offer a more concise and interpretable representation of the data. The optimal value for the hyperparameter $\alpha_L$ is chosen using cross-validation, and the resulting non-zero coefficients in $\theta$ can be interpreted as the most critical features for immunomodulation, as shown in Figure S4.

The linear nature of Equation S7 makes it easy to interpret the sign of the learned coefficients: large negative weights $\theta_k < 0$ indicate features that are negatively correlated with immunomodulation, while large positive weights $\theta_k > 0$ indicate features that are positively correlated with immunomodulation. As shown in Figure S5, by examining the rank-ordering of the coefficients with the largest absolute values, we can identify structural fragments that are most informative for predicting immunomodulation. There were, in total, 33 chemical fragment descriptors that were retained by at least one of the LASSO models for different immunological objectives. In addition, our trained LASSO model can be used to predict immunomodulation for larger molecules and/or molecules not contained within the training data. A CSV file showing the code names and the corresponding chemical interpretations of these 33 chemical fragment descriptors is provided in the Supporting Information.

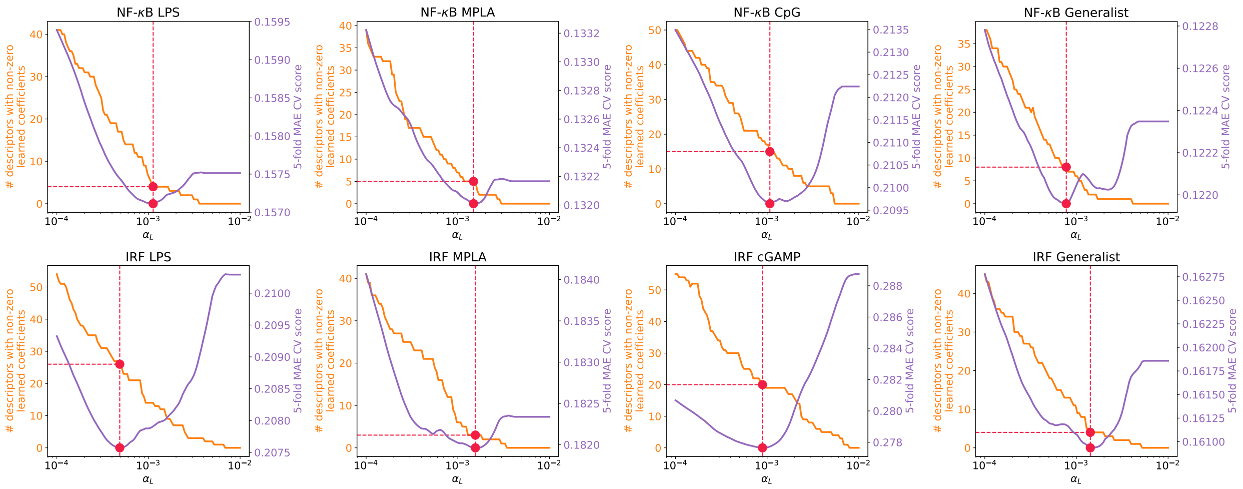

Figure S4: **The performance of LASSO regression models evaluated with respect to sparsity regularization parameter $\alpha_L$.** The plot presents the number of molecular descriptors that have non-zero learned coefficient values, which were identified by training the model using a range of $\alpha_L$ values (displayed on the left y-axis in orange). Additionally, the plot shows the 5-fold cross-validation mean absolute error (MAE) score of the LASSO regression model at each corresponding $\alpha_L$ value (displayed on the right y-axis in purple). The optimal $\alpha_L$ is shown in red points where the corresponding MAE is the lowest. The title of each plot denotes the corresponding immune signaling pathway and agonist used for stimulation.

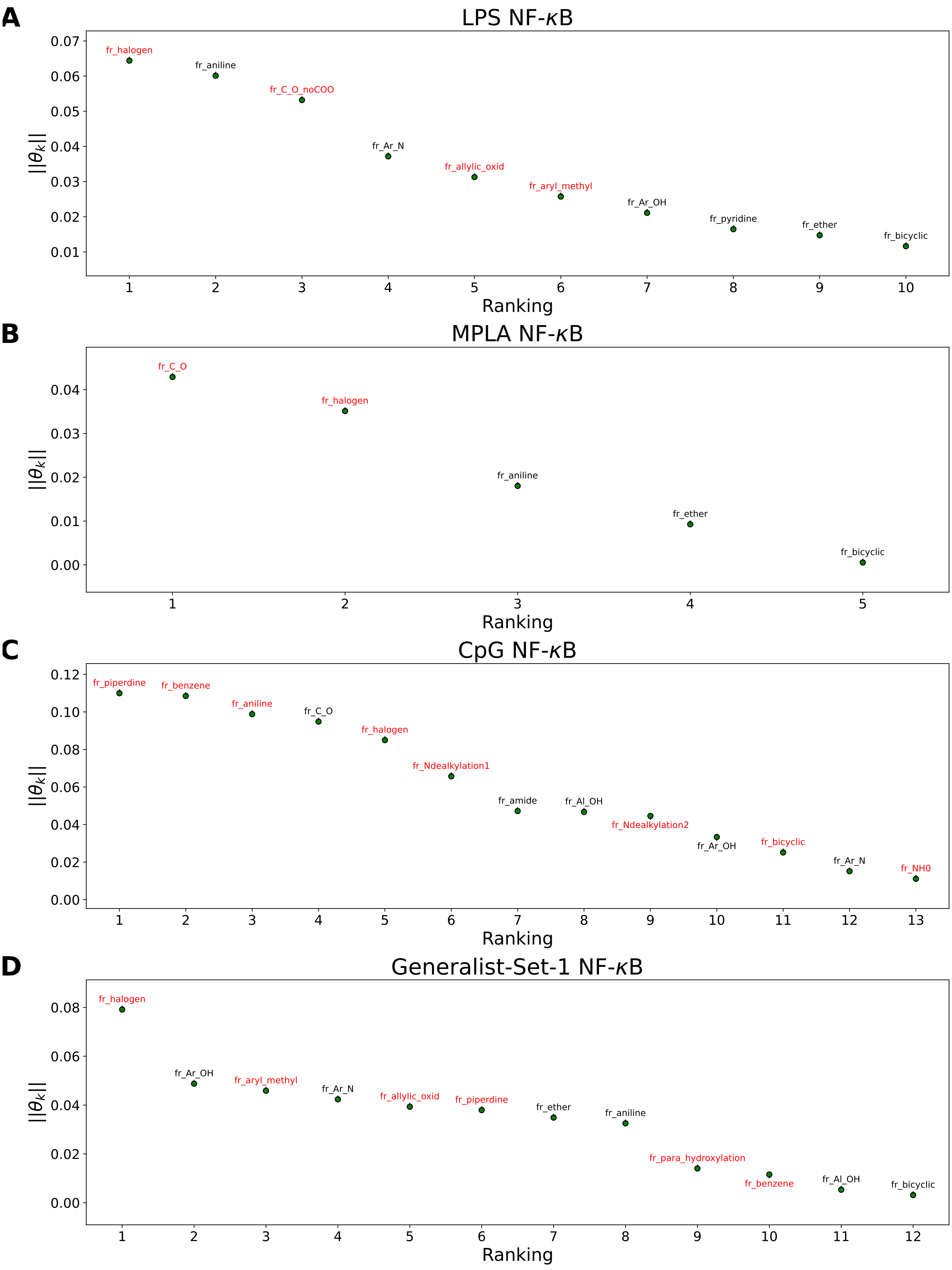
A
LPS NF-κB
fr_halogen
fr_aniline
fr_C_O_noCOO
fr_Ar_N
fr_allylic_oxid
fr_aryl_methyl
fr_Ar_OH
fr_pyridine
fr_ether
fr_bicyclic
||θ_k||
Ranking
B
MPLA NF-κB
fr_C_O
fr_halogen
fr_aniline
fr_ether
fr_bicyclic
||θ_k||
Ranking
C
CpG NF-κB
fr_piperdine
fr_benzene
fr_aniline
fr_C_O
fr_halogen
fr_Ndealkylation1
fr_amide
fr_Al_OH
fr_Ndealkylation2
fr_Ar_OH
fr_bicyclic
fr_Ar_N
fr_NH0
||θ_k||
Ranking
D
Generalist-Set-1 NF-κB
fr_halogen
fr_Ar_OH
fr_aryl_methyl
fr_Ar_N
fr_allylic_oxid
fr_piperdine
fr_ether
fr_aniline
fr_para_hydroxylation
fr_benzene
fr_Al_OH
fr_bicyclic
||θ_k||
Ranking

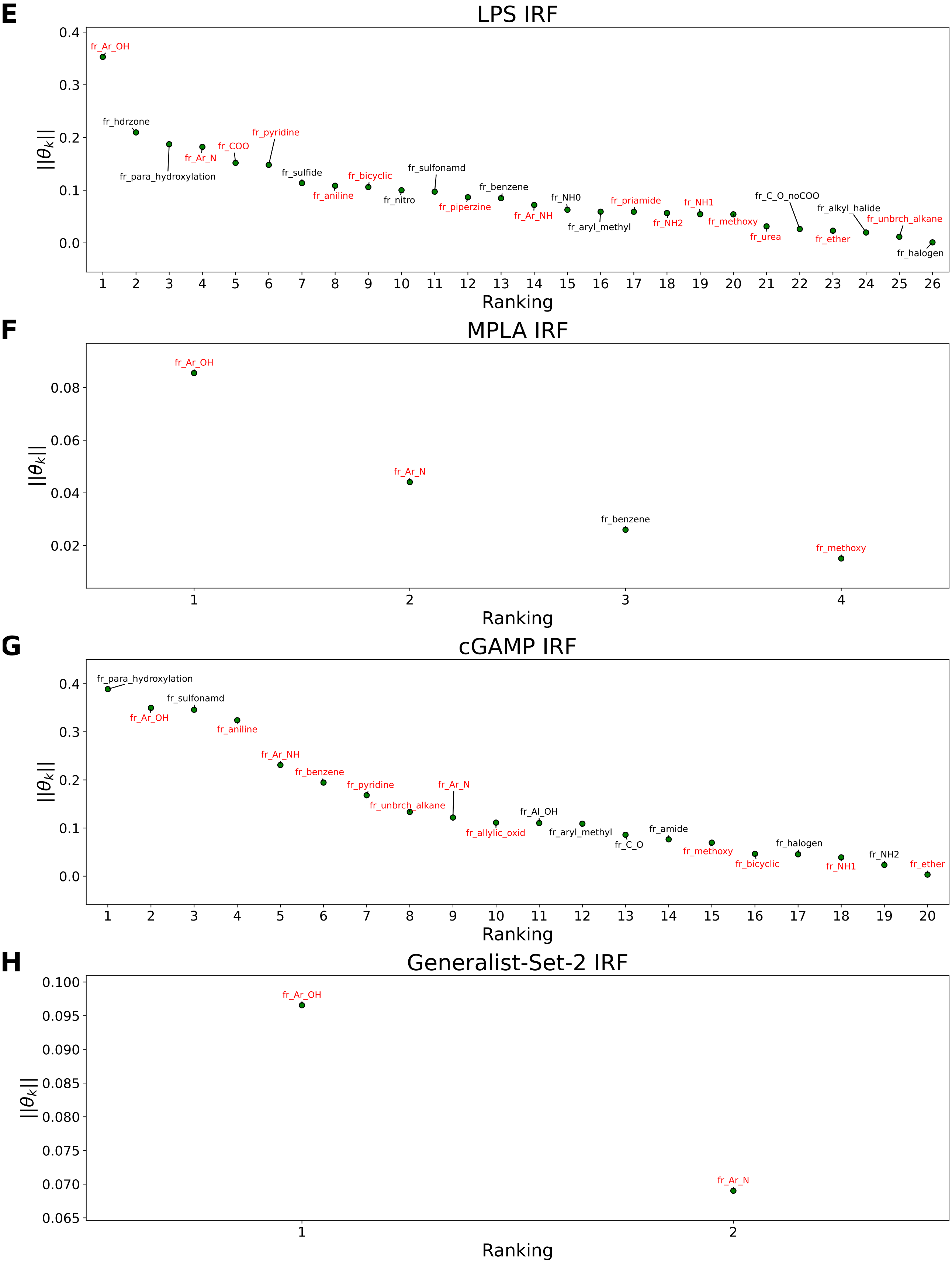
E
LPS IRF
||$\theta_k$||
fr_Ar_OH
fr_hdrzone
fr_para_hydroxylation
fr_Ar_N
fr_COO
fr_pyridine
fr_sulfide
fr_aniline
fr_bicyclic
fr_nitro
fr_sulfonamd
fr_piperzine
fr_benzene
fr_Ar_NH
fr_NH0
fr_aryl_methyl
fr_priamide
fr_NH2
fr_NH1
fr_methoxy
fr_urea
fr_C_O_noCOO
fr_ether
fr_alkyl_halide
fr_unbrch_alkane
fr_halogen
Ranking
F
MPLA IRF
||$\theta_k$||
fr_Ar_OH
fr_Ar_N
fr_benzene
fr_methoxy
Ranking
G
cGAMP IRF
||$\theta_k$||
fr_para_hydroxylation
fr_Ar_OH
fr_sulfonamd
fr_aniline
fr_Ar_NH
fr_benzene
fr_pyridine
fr_unbrch_alkane
fr_Ar_N
fr_allylic_oxid
fr_Al_OH
fr_aryl_methyl
fr_C_O
fr_amide
fr_methoxy
fr_bicyclic
fr_halogen
fr_NH1
fr_NH2
fr_ether
Ranking
H
Generalist-Set-2 IRF
||$\theta_k$||
fr_Ar_OH
fr_Ar_N
Ranking

Figure S5: **Full accounting of non-zero learned coefficient weights $\theta_k$ associated with substructure features ranked by their magnitudes of weights in units of log2-fold change.** For each molecule $n$, we generate a standardized feature vector over all 70 features $k$, and collect these within a feature matrix $F_{n,k}$. Using this feature matrix, we apply LASSO regression to predict the calculated immunomodulation acquired from high throughput screening experiments and identifying the learned non-zero coefficient weights $\theta_k$ with the highest magnitude from the reduced feature set retained by the LASSO model. The features with positive weights are displayed in black text, while the ones with negative weights are displayed in red text. Positive weights imply that there is a positive correlation between the feature values and enhanced immunomodulation, while negative weights indicate a positive correlation between the feature values and inhibitory immunomodulation.

### S2 Supplementary Experimental Methods and Results

#### S2.1 High throughput screening experiments

##### S2.1.1 Immonomodulators delivered alone do not affect immune responses of NF-$\kappa$B and IRF pathways

The raw data captured by the plate reader in our high throughput screening (HTS) experiments, which reflects the magnitude of the immune response induced by each agonist or the combination of an agonist and an immunomodulator, is presented in Figure S6. LPS, MPLA and CpG are generally good NF-$\kappa$B agonists while cGAMP struggles in activating NF-$\kappa$B pathway. LPS, MPLA and cGAMP are typically effective in activating the IRF pathway, while CpG faces difficulties in initiating the IRF pathway. Most candidate immunomodulators do not stimulate an immune response on their own, without the presence of agonists. There were only 27/2880 (=0.9%) immunomodualtor candidates with which the samples show absorbance $> 0.5$.

#### S2.2 Low throughput immunomodulation validation experiments

##### S2.2.1 Assay used for cytokine release profile testing

Monocytes were harvested from 6-week-old C57BL/6 mice and were differentiated into dendritic cells (BMDCs) using supplemented culture medium: RPMI 1640 (Life Technologies), 10% HIFBS (Sigma-Aldrich), Recombinant Mouse GM-CSF (carrier-free)(20 ng/ml; BioLegend), 2 mM l-glutamine (Life Technologies), 1% antibiotic- antimycotic (Life Technologies), and 50$\mu$M $\beta$-mercaptoethanol (Sigma-Aldrich). After 6 days of culture, BMDCs were plated at 100,000 cells per well and incubated with modulator (10 $\mu$M). After 1 hour, agonist was added. Cells were incubated for 24 hours at 37°C and 5% CO2. Supernatant cytokines were measured using Legendplex Mouse Inflammation Cytokine Kit (Biolegends) or a VeriKine IFN-$\beta$ ELISA (PBL Assay Science).

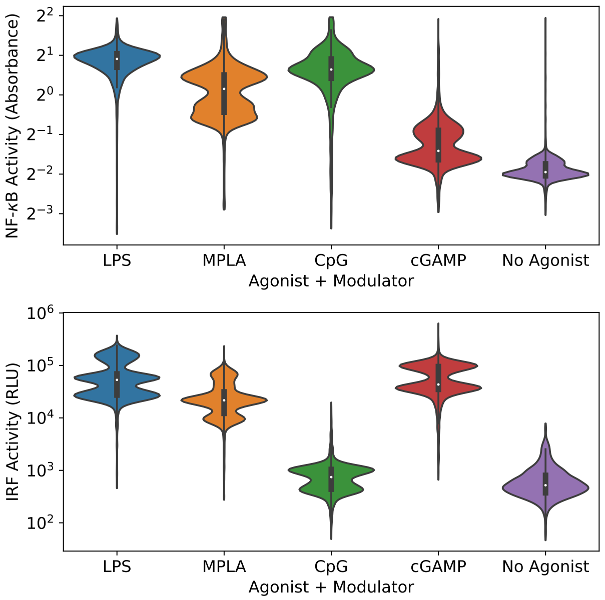

Figure S6: **Results of active learning screen as raw data.** NF-$\kappa$B activity was measured as absorbance readings at 620 nm, and IRF activity was measured as luminescence readings in the units of relative light unit (RLU). The higher the absorbance or luminescence, the stronger the corresponding immune activity. The CpG agonist shows minimal stimulation of the IRF pathway relative to no agonist present and was dropped from subsequent analyses.

##### S2.2.2 Immonomodulators delivered alone do not affect cytokine production

For transcription factor activity, modulators alone do not influence the immune responses (Figure S6). This also applies to cytokine secretion profile in that modulator do not stimulate cytokine release in the absence of agonist stimulation (Figure S7). It is the combination of agonist and immunomodulators that allow for significant enhancement and inhibition of both transcription factor activity and cytokine secretion.

##### S2.2.3 Comparing PME-4007 and MSA-2

MSA-2 is a recently discovered potent STING agonist that can stimulate STING pathway and was identified as a chemical inducer of interferon-$\beta$ secretion[30] via a high throughput screening process which involved ~2.4M compounds. We subjected our leading IFN-$\beta$ inducing compound, PME-4007, to additional comparisons of its cytokine profile in the presence of cGAMP to that of MSA-2. We largely followed the protocol mentioned in Section 2.3

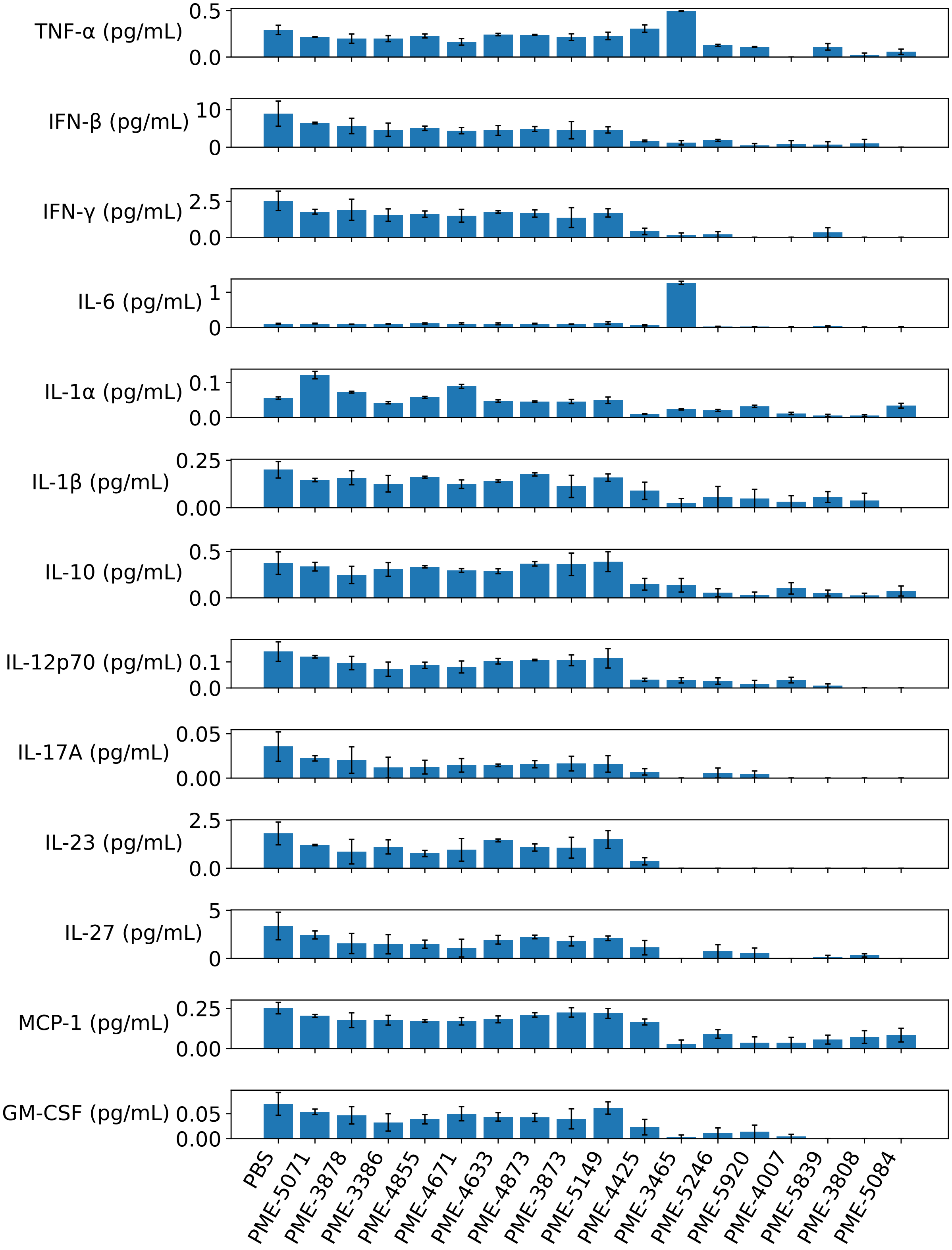
TNF-α (pg/mL)
IFN-β (pg/mL)
IFN-γ (pg/mL)
IL-6 (pg/mL)
IL-1α (pg/mL)
IL-1β (pg/mL)
IL-10 (pg/mL)
IL-12p70 (pg/mL)
IL-17A (pg/mL)
IL-23 (pg/mL)
IL-27 (pg/mL)
MCP-1 (pg/mL)
GM-CSF (pg/mL)
PBS
PME-5071
PME-3878
PME-3386
PME-4855
PME-4671
PME-4633
PME-4873
PME-3873
PME-5149
PME-4425
PME-3465
PME-5246
PME-5920
PME-4007
PME-5839
PME-3808
PME-5084

Figure S7: **Modulators (10 $\mu$M) alone minimally affects cytokine production 24 hours after addition.** Each panel shows the secreted concentration for a specific cytokine stimulated by PBS (as negative control) and 17 selected top-performing immunomodulator candidates (without the addition of agonists), addressed for 13 cytokines. The cytokine production result from the addition of modulators in the absence of agonists is within the same order of magnitude as that of the PBS negative control, showing that the addition of immunomodulators alone minimally affects the production of cytokines. Although for IL-6 production, there is one molecule PME-3465 that has much higher IL-6 secretion than PBS control, the magnitude of the amount of IL-6 being released is still minimal immunologically. The cytokine production is measured by LegendPlex.

in the main text. To obtain more stringent results for this low throughput comparison, we used more replicates ($N$=6). Considering the moderate cytotoxicity exhibited by PME-4007, we used a lower concentration at 2 $\mu$M instead of the generic concentration of 10 $\mu$M for all immunomodulators in all preceding experiments. To measure the cell viability of this experiment, we utilized the MTT assay. After BMDCs were incubated with or without the immunomodulator PME-4007 and with the agonist cGAMP or MSA-2 overnight, the cell supernatant was removed and 70 $\mu$L of PBS was added. Cells were incubated with 10 $\mu$L of 5 mg/mL (4,5-dimethylthiazol-2-yl)-2,5-diphenyltetrazolium bromide (MTT) for 3 hours at 37°C. The medium was then removed, and 70 $\mu$L of DMSO was added to solubilize formazan crystals. Plates were then read at 540 nm using a BioTek plate reader. Cell viability was calculated as a percentage of proliferation versus PBS-treated negative control cells. The results of MTT assay are shown in Figure S8A. Figure S8B and Figure S8C, as extensions to Figure 6B in the main text, show that the modulation profile of PME-4007 over cytokine secretion is not limited to IFN-$\beta$ and TNF-$\alpha$.

#### S2.3 Safety statement

During all wet lab experiments involved in this work, there were no unexpected or unusually high safety hazards encountered.

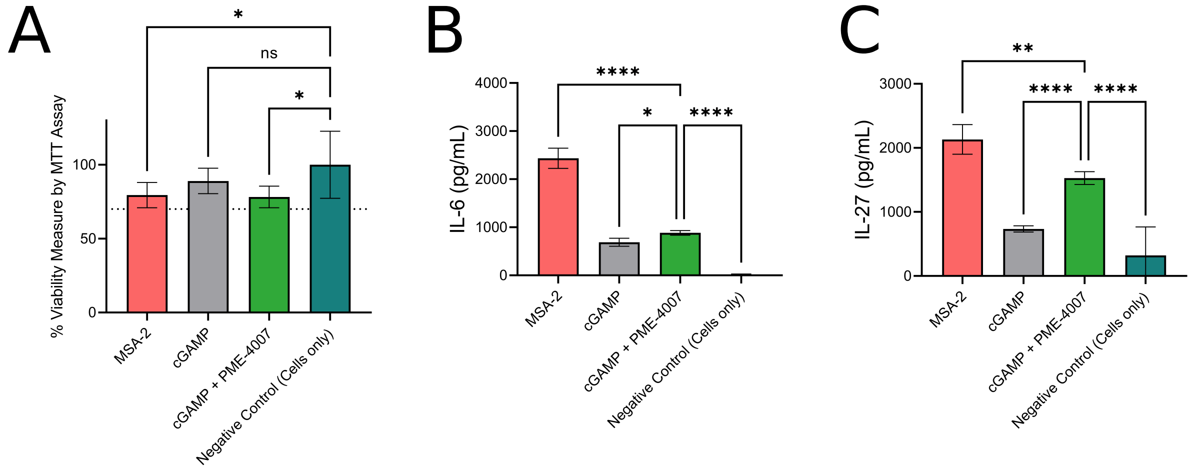

Figure S8: **PME-4007 induces minimal cytotoxicity while enhancing the secretion of IL-6 and IL-27 stimulated by cGAMP.** (A) MTT Assay indicates that MSA-2, cGAMP, and the combination of cGAMP and PME-4007 induce minimal cytotoxicity to the cells as the cell viability is all higher than the cutoff 70%, and close to the level of negative control resting cells. (B) PME-4007 slightly enhances the secretion of IL-6. MSA-2 strongly stimulates the secretion of IL-6 and is much stronger than the stimulation induced by cGAMP. (C) PME-4007 significantly enhances the secretion of IL-27 with cGAMP, and it is close to the level of stimulation induced by MSA-2.

#### S3 Supplementary Figures

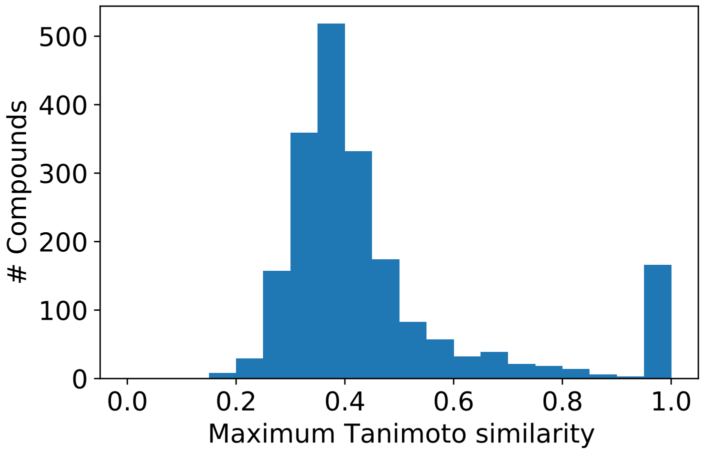

Figure S9: **Maximum pairwise Tanimoto similarities between the 2674 previously screened molecules used to train our initial GPR models and the top-performing candidates capable of >2-fold enhancement/inhibition identified by our active learning screen.** Tanimoto similarities between all pairs associating one molecule in the previous screen and one molecule in the active learning screen were computed with ECFP4 molecular fingerprint using RDKit.[31] The maximum Tanimoto similarities for each of the molecules identified in the active learning screen were determined by $\max_j \text{Tanimoto}(\mathbf{x}_i, \mathbf{x}_j)$, where $\mathbf{x}_i$ denoted the $i^{\text{th}}$ molecule in the active learning screen and $\mathbf{x}_j$ denoted the $j^{\text{th}}$ molecule in the previous screen. The Tanimoto similarity quantifies the proportion of chemical substructures shared by a pair of molecules and it is a continuous number between 0 and 1.0, where 1.0 denotes complete identity. The peak near 1.0 indicates that the model samples molecules with similar structures to known good immunomodulators, but the large support for the histogram in the 0-0.6 region indicates that the active learning screen is also exploring molecules that are substantially diverse from the initial training data.

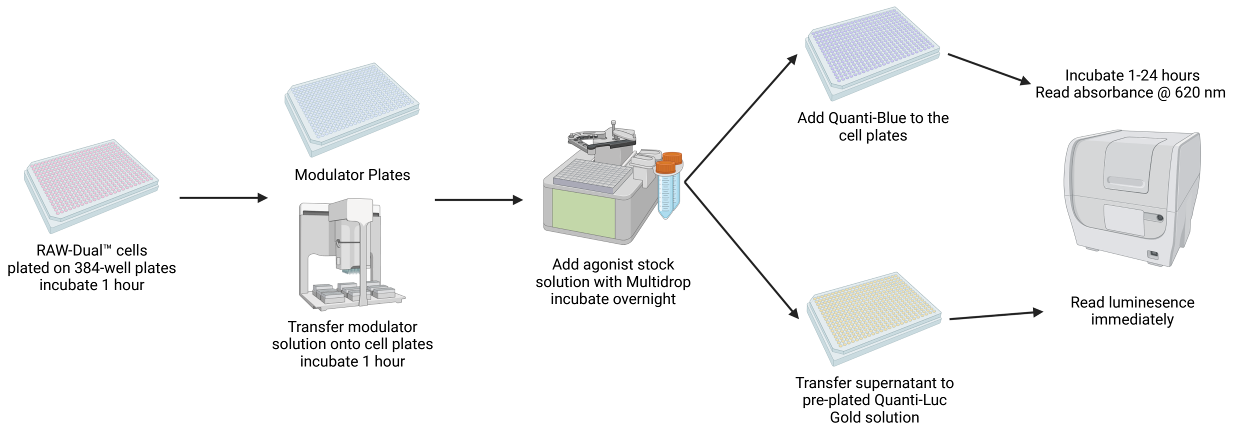

Figure S10: **High-throughput experimental screening protocol for NF-$\kappa$B transcription factor levels and IRF activity.** We seeded 50,000 RAW-Dual™ macrophages in 384-well plates in 45 $\mu$L of complete media using a MultiDrop™ Combi liquid handler, then incubated for 1 hour at 37°C. We transferred immunomodulator compounds from source plates (10mM dissolved in dimethyl sulfoxide, DMSO) to a final concentration of 10 $\mu$M using a Janus® G3 via pintool, then incubated for another 1 hour at 37°C. One of the four PRR agonists was added in 5 $\mu$L of media to achieve the desired concentration into corresponding wells using a MultiDrop™ Combi liquid handler. For control plates where we test the effect of modulators in the absence of agonists, we added same volume of media instead. We incubated the cells overnight. We then add QUANTI-Blue™ and QUANTI-Luc™ reagents as appropriate, and read the absorbance at 620 nm and the luminescence using a BioTek Synergy™ Neo2 Hybrid multimode microplate reader, which correspond to the NF-$\kappa$B transcription factor levels and the IRF activity levels, after necessary calibration and standardization. Portions of this figure were created with `BioRender.com`.

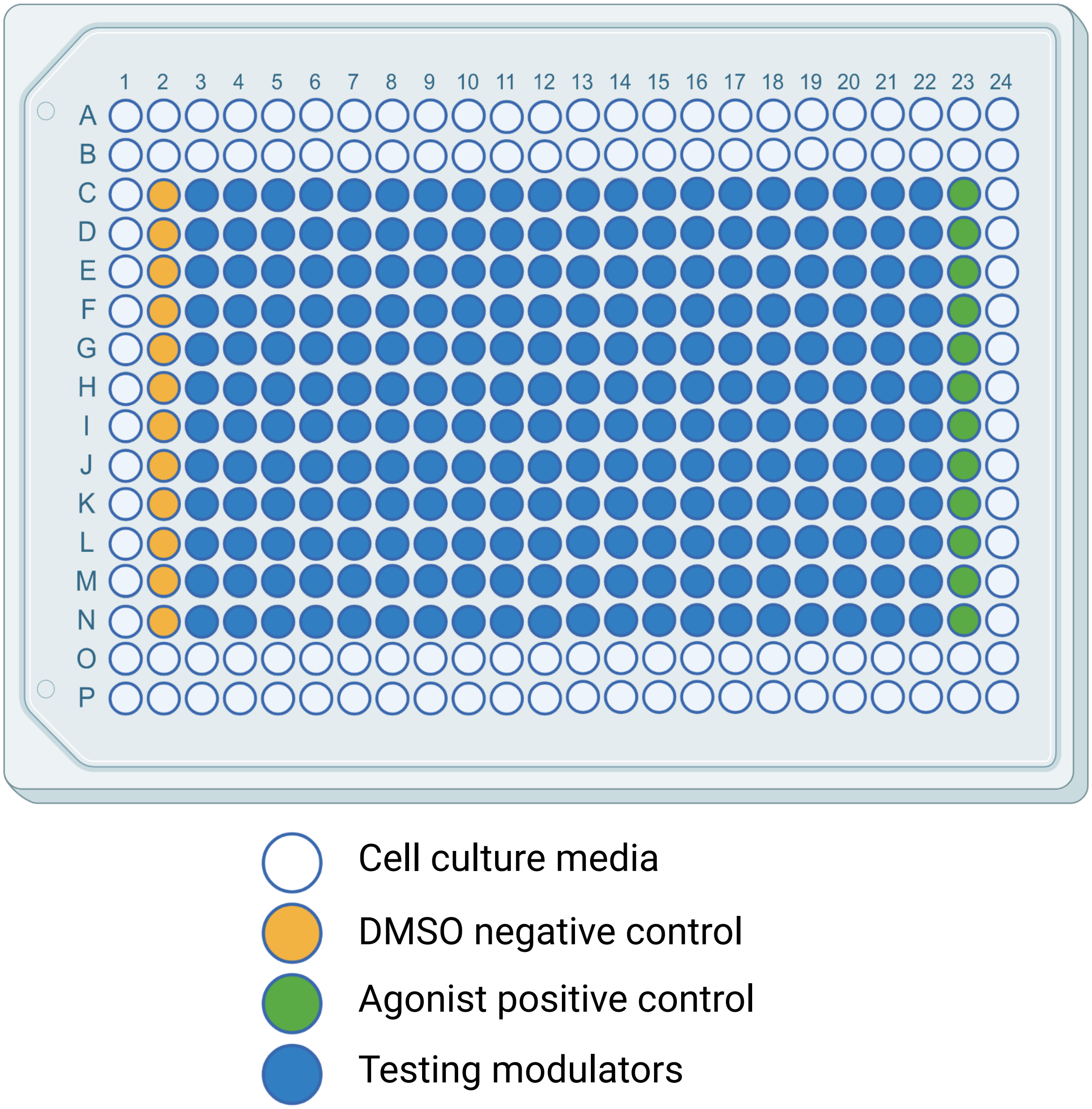

Figure S11: **384-well plate layout.** One column and two rows on edges are not used to test and filled with cell culture media to avoid error induced by evaporation of water. Portions of this figure were created with `BioRender.com`.

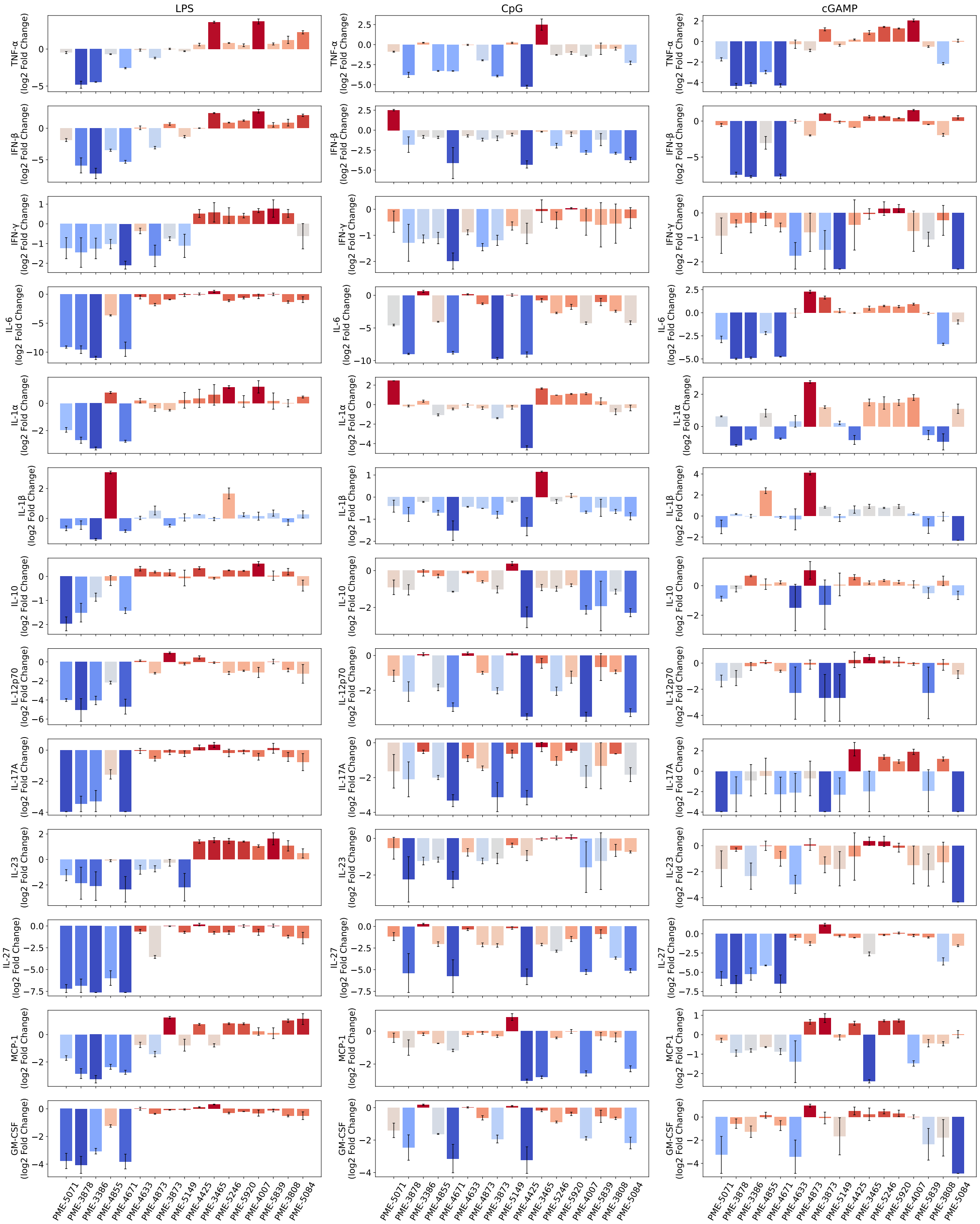

Figure S12: **Immunomodulation of 17 selected top-performing candidates over cytokine release profiles activated by LPS, CpG and cGAMP.** The extent of immunomodulation is visualized as bar plots with error bars indicating standard errors associated with each experiment. The bars are also color-coded by the log2-fold change values for better clarity in showing those immunomodulation with large extent, representing potent enhancers or suppressors.

### S4 Supplementary Tables

Table S5: **Source library of screened molecules.** The table shows (1) the number of compounds in each chemical screening library, (2) the number of compounds experimented with HTS in each library, (3) the number of good modulators (with >2-fold modulation) found in each library and (4) the ratio of good modulators (the 3$^{rd}$ column divided by the 1$^{st}$ column). The last three libraries, namely Microsource Spectrum Collection, Prestwick Chemical Library and Selleckchem FDA-approved Drug Library, possess relatively higher ratios of good modulators identified. The statistics excluded non-viable compounds.

| Chemical Screening Library | # compounds | # experimented | # good modulators | % good modulators |
|---|---|---|---|---|
| ChemBridge DS550-3 | 29547 | 571 | 52 | 0.18 % |
| ChemBridge ES550-2 | 49584 | 657 | 54 | 0.11 % |
| Life Chemicals Biologically Active Compound Library | 7938 | 176 | 27 | 0.34 % |
| Life Chemicals Pre-plated Diversity Sets PS4 | 50233 | 546 | 22 | 0.04 % |
| Micro Source The Spectrum Collection | 1821 | 350 | 10 | 0.55 % |
| Prestwick Chemical Library | 1219 | 276 | 9 | 0.74 % |
| Selleckchem FDA-approved Drug Library | 1499 | 708 | 26 | 1.73 % |

Table S6: **List of agonists studied and their working concentration.**

| Agonist | Target | Concentration ($\mu$g/mL) | Reference |
|---|---|---|---|
| LPS | TLR4 | 0.1 | [32] |
| MPLA | TLR4 | 0.25 | [33] |
| CpG (ODN 1826) | TLR9 | 0.5 | [34] |
| cGAMP | STING | 10 | [22] |
